# Decomposing predictability to identify dominant causal drivers in complex ecosystems

**DOI:** 10.1101/2022.03.14.484197

**Authors:** Kenta Suzuki, Shin-ichiro S. Matsuzaki, Hiroshi Masuya

## Abstract

Ecosystems are complex systems of various physical, biological, and chemical processes. Since ecosystem dynamics are composed of a mixture of different levels of stochasticity and nonlinearity, handling these data is a challenge for existing methods of time-series based causal inferences. Here we show that, by harnessing contemporary machine learning approaches, the concept of Granger causality can be effectively extended to the analysis of complex ecosystem time series and bridge the gap between dynamical and statistical approaches. The central idea is to use an ensemble of fast and highly predictive artificial neural networks to select a minimal set of variables that maximizes the prediction of a given variable. It enables decomposition of the relationship among variables through quantifying the contribution of an individual variable to the overall predictive performance. We show how our approach, EcohNet, can improve interaction network inference for a mesocosm experiment and simulated ecosystems. The application of the method to a long-term lake monitoring dataset yielded new but interpretable results on the drivers causing cyanobacteria blooms, which is a serious threat to ecological integrity and ecosystem services. Since performance of EcohNet is enhanced by its predictive capabilities, it also provides an optimized forecasting of overall components in ecosystems. EcohNet could be used to analyze complex and hybrid multivariate time series in many scientific areas not limited to ecosystems.

**Significance Statement:** Effective use of ecosystem monitoring data to resolve global environmental issues is a major challenge of the 21st century ecology. A promising solution to address this challenge is a time-series-based causal analysis which can provide insight on the mechanical links between ecosystem components. In this work, a model-free framework named EcohNet is proposed. EcohNet utilizes ensemble predictions of echo state networks, which are known to be fast, accurate, and highly relevant for a variety of dynamical systems, and can robustly predict causal networks of ecosystem components. It also can provide an optimized forecasting of overall ecosystem components, and could be used to analyze complex and hybrid multivariate time series in many scientific areas, not limited to ecosystems.

## Introduction

Various systems in the world, from cells, organisms, ecosystems, and our own societies, are complex and driven by many interacting components. An attempt to infer relationships among the components as a causal network is an important step in understanding their mechanistic basis. Great efforts have been made to infer such networks from time series [1, 2]. In particular, methods that can overcome the limitations of classical methods [3, 4], such as CCM [5], which is suitable for systems under the influence of dynamic processes, and PCMCI [6], which is suitable for stochastic systems, have been proposed and are becoming widely used.

However, there is a problem when attempting to apply the existing time-series-based causal analyses to ecosystems. Ecosystems are complex systems of various physical, biological, and chemical processes [7]. Meteorological variables such as temperature and precipitation are under the influence of atmospheric and oceanic fluid dynamics [8]. Recent studies showed that these earth system dynamics are well captured as stochastic processes, and approaches based on statistical causal analysis are effective in elucidating their relationships [2, 6, 9]. In contrast, the density and abundance of organisms often show strong nonlinear dynamics, driven by interactions between organisms. This motivates the application and success of approaches based on nonlinear dynamical systems [5, 10-12]. Some chemicals control ecological dynamics as essential resources and are produced by organismal activities, as well they are under the control of global geochemical cycles [13]. These processes influence each other, but it is through rather weak causal couplings, and the dynamics of any one process may not become dominant [5, 14-17]. Methods for revealing causal relationships among ecosystem components from time series data need to be robust to dynamical complexity, i.e., the different levels of stochasticity and nonlinearity. However, the assumptions of most methods do not satisfy this requirement [1, 18].

In this paper, we introduce a new method called EcohNet. It is based on the ensemble prediction of neural networks [19-21] that can seamlessly handle stochastic/deterministic and linear/nonlinear dynamics [22]. Therefore, it is expected to be robust to dynamical complexity. The ensemble prediction is used to decompose relationships among variables in terms of predictability [23]. Here, the contribution of one variable to predictive performance can be evaluated apart from that of the other variables, as in the concepts of partial correlation and conditional independence. It is expected that even if some variables are driven by a strong driver, weak relationships among variables can be detected separately from the effect of the driver without special treatments [24-27].

Although various methods have been proposed and used for descriptive purposes, the relevance of their application to actual ecosystem monitoring data has not been seriously examined. How well can interactions between ecosystem components be captured as causal relationships in EcohNet? What advantages does it have over conventional methods used to assess causal analysis? To answer these questions, we performed benchmarking with data from long-term observation of an aquatic mesocosm, as well as a simulated dataset to test robustness to different dynamical complexity (equilibrium, equilibrium forced by an external oscillator, and intrinsic oscillatory dynamics, as well as different magnitudes of noise) under two interaction types (food web and random interaction) and three observational conditions (different data size, sampling interval and the presence of unobserved species). In addition to the performance criteria for network inference, we focused on susceptibility to the often-problematic interaction topologies such as chain relationships (e.g., in a three-species food chain X←Y←Z, a causal relationship may be identified between X and Z) and fan-out relationships (e.g., in a one-predator-two-prey-relationships X←Z→Y, a causal relationship may be identified between X and Y) [5, 24, 28]. We then applied our method to a long-term lake monitoring dataset that includes heterogeneous components such as meteorological, chemical, and biological variables, and interpreted the results. Here, we mainly focused on how EcohNet provides insight on the drivers of cyanobacteria blooms, which is often a serious threat to ecological integrity and ecosystem services [29], but also we show how well the detection of unrealistic causal relationships such as those from biological to meteorological variables were avoided.

### EcohNet

EcohNet combines a type of recurrent neural network, called echo state network (ESN) [19-20, 30], with a progressive selection of variables [31] (Figure 1; see Materials and Methods for detail). Initially, a target variable is selected (Fig. 1A). Then, *N* ESNs are generated and the ensemble predictive performance (prediction skill) of one-time-step ahead is evaluated when the past state of the target itself is given as an input of ESNs (Figure 1B,C). As the result, we obtain a distribution of prediction skills (illustrated by a gray mountain shape in Fig.1B). In the next step, we choose a second variable other than the target variable one by one and adopt the one that improves the prediction skill the most. Here, *N* ESNs are generated for each pair of variables. If the prediction skill does not improve, no further new variables are adopted. This process is repeated for variables that have not yet been selected, as long as new variables are adopted. As a result of the above process, a set of variables 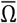 that maximizes the prediction skill for the target variable is obtained. Then, 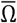 is used to evaluate the *unique prediction skill*, which represents the unique contribution of each variable in 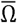 to the overall prediction skill (Figure 1D). If 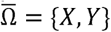 and the target variable is *X*, for each variable (here, *X* and *Y*), we evaluate how much the prediction changes when one variable is excluded from it (e.g., 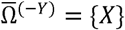). For example, when denoting the prediction skill for 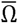 and 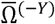as, 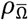and 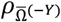 respectively, the unique contribution of Y on X is 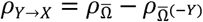. In this stage, a variable is removed from 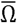 if 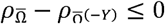. By performing the above procedure with all variables as targets in turn, a prediction-skill-based causal network is obtained for the entire system.

**Figure 1.**
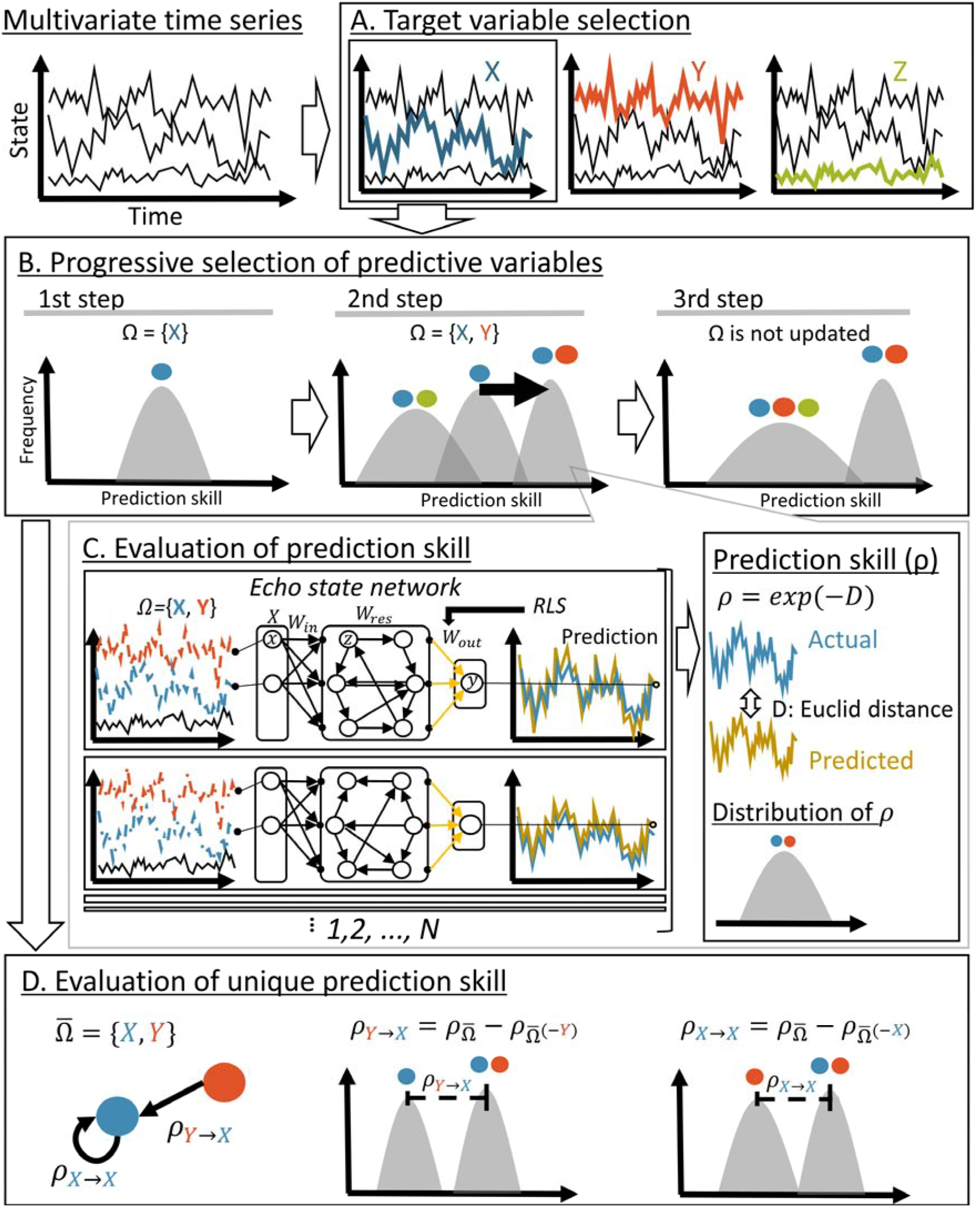
Graphical explanation of EcohNet. See main body and Materials and Methods for the explanation.

To show how EcohNet works, we explain what would be 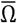 for X in some representative relationships of the three variables X, Y and Z. First, we assume a transitive relationship (Z→Y→ X and Z→X; SI Appendix Fig. S1A). Here, in addition to *ρ*_*X*→*X*_, both *ρ*_*Y*→*X*_and *ρ*_*Z*→*X*_ are non-zero corresponding to the direct relationship Y→X and Z→X. In a chain relationship (Z→Y→X; SI Appendix Fig. S1B) the direct relationship Z→X is removed. In this case, the contribution of Z on X is always mediated by Y. Thus, if all else is equal, only *ρ*_*Y*→*X*_and *ρ*_*X*→*X*_are expected to have non-zero values. In a fan-out relationship (X←Z→Y; SI Appendix Fig. S1C) the direct relationship Y→X is removed. In this case, both X and Y have a direct relationship with Z but remain disconnected from each other. X and Y share dynamic imprint of Z that may contribute to an ability to predict X from Y but it is simply involved in the direct influence of X on Z. Thus, if all else is equal, only *ρ*_*X*→*X*_and *ρ*_*Z*→*X*_are expected to have non-zero value. The importance of evaluating the unique prediction skill is highlighted by another representative example (SI Appendix Fig. S2). In this case, both Z_1_ and Z_2_ have a fan-out relationship with X and Y (X←Z_1_→Y, and X←Z_2_→Y). As explained in Runge et al. [1], a simple forward stepwise algorithm would select Y first followed by Z_1_ and Z_2_, because Y can have a larger individual contribution than either Z_1_ or Z_2_. However, evaluation of the unique prediction skill can remove Y from 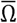 since contribution of Y is included in the union of the contribution of Z_1_ and Z_2_, and thus 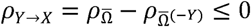.

### Conventional approaches

We selected a correlation-based method (Spearman rank correlation), two equation-free methods from nonlinear time series analysis (convergent cross mapping (CCM) [5] and partial cross mapping (PCM) [28]) and an equation-based method (LIMITS) [32] as the conventional approaches for comparison with EcohNet (please see SI Appendix A for the details of the implementations), and compared them using the evaluation criteria of a binary classification task (SI Appendix Fig. S3). CCM and PCM are intended to detect causality in dynamical system and are therefore comparative to EcohNet. Since there are several different implementations for CCM, we tested three representative approaches (SI Appendix A2). PCM is an extension of CCM that accounts for direct causality between two variables conditional on indirect causation through a third variable. Thus, PCM implements an idea similar to the unique prediction skill of EcohNet. LIMITS infers the strength of ecological interactions directly rather than the causal relationships among variables. In the benchmark using simulation data, the results of LIMITS should be carefully interpreted because there is the advantage that the basic process is consistent, i.e., it assumes Lotka-Volterra (LV) equation used to generate the data. In other words, it contains more prior knowledge than the other methods. Applying it to relationships of all ecosystem components such as chemical and meteorological variables exceeds its scope of application. Although it is frequently pointed out that correlation does not signal the presence of interaction, we considered Spearman rank correlation as a baseline.

## Results

### Benchmarking

When applied to the data from a long-term mesocosm experiment [33], EcohNet identified 11 out of 13 interacting pairs of components, with 3 false positives, and was superior to conventional methods (Fig. 2a-d). It outperformed other methods in all evaluation criteria (Fig. 2e). The results were not sensitive to different parameter values that might affect the performance of echo state networks, except when the forgetting factor *λ* was very close to 1 (see SI Appendix B). A benchmark with datasets generated by food web models (SI Appendix Fig. S4) also supported the superiority of our method. ROC-AUC and F1-score of EcohNet outperformed CCM, PCM and Spearman rank correlation except for ROC-AUC of CCM in *O*_*s,L*_ in which the signal of interaction was considered to be strong (Fig. 2g,h). Performance of CCM was largely reduced in *E*_*s,L*_and *EO*_*s,L*_. PCM performed better than CCM in *E*_*s,L*_, but it was reduced in *EO*_*s,L*_. CCM was better than PCM in *EO*_*s,L*_potentially because of the use of seasonal surrogate method. Performance of Spearman rank correlation was comparable to other methods only in *E*_*s,L*_. LIMITS outperformed EcohNet in *O*_*s,L*_, especially in ROC-AUC. Its high applicability to cases of intrinsic oscillations has been reported in previous studies [34]. It is also worth noting that EcohNet’s performance was comparable to LIMITS in most cases, despite not explicitly assuming underlying LV processes.

**Figure 2.**
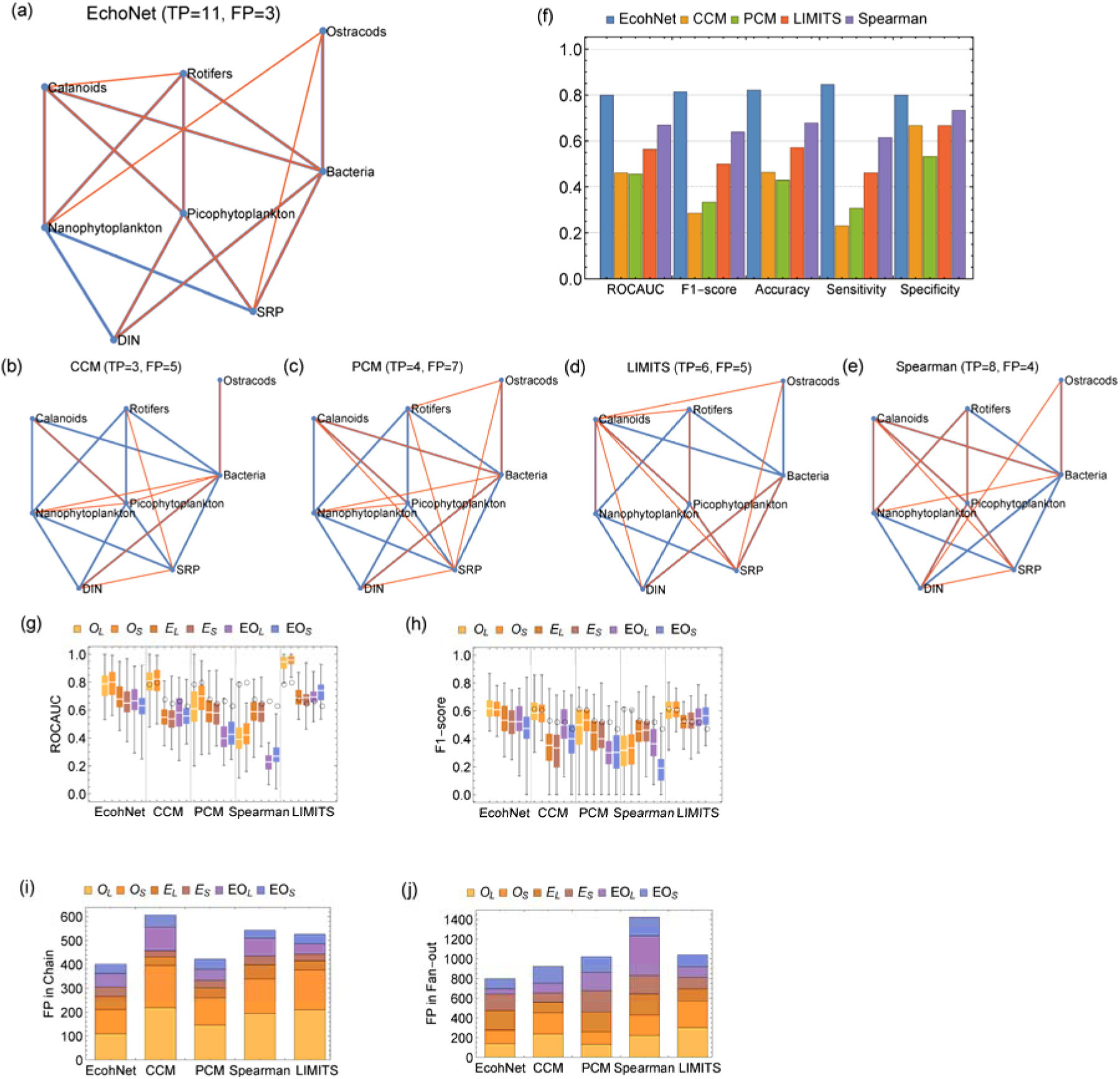
Benchmarking results. (a-e) Actual (blue) and inferred (red) network of EcohNet and conventional approaches (CCM, PCM, Spearman rank correlation and LIMITS) for mesocosm data (DIN and SRP stands for total dissolved inorganic nitrogen and soluble reactive phosphorus, respectively). The number of true positives (TP) and false positives (FP) are shown above the panel. (f)The value of evaluation criteria for mesocosm data. (g, h) Two performance criteria (ROC-AUC and F1-Score) for the food web models. Here, O is intrinsic oscillation, E is equilibrium and EO is equilibrium forced by an external oscillator and the suffixes S and L indicate small and large noise magnitude, respectively (see SI Appendix D2 for simulation parameter values). In the box plot, white lines indicate the median, box edges indicate the first and third quartile value, and whiskers indicate maximum and minimum values. Black circles indicate the median value of EcohNet. In (g) and (h), the result of LIMITS should be carefully interpreted because it assumes Lotka-Volterra (LV) equation used to generate the benchmark data. (i) Cumulative number of incorrectly identified links in chain relationships (for 6×100=600 time series in total). (j) Cumulative number of incorrectly identified links in fan-out relationships (for 6×100=600 time series in total).

Although accuracy of EcohNet, CCM and PCM was consistent among *O*_*s,L*_, *E*_*s,L*_and *EO*_*s,L*_except for PCM for *EO*_*s,L*_, there were large variations in sensitivity and specificity for CCM and PCM (Fig. 3a-c). The number of detected links varied depending on the simulation conditions (Fig. 3d). It suggests that the fundamental sensitivity of the methods used to detect presence of interactions is affected by dynamic complexity. In comparison, performance of EcohNet was relatively stable, and its sensitivity, specificity and the number of links detected were not greatly affected by the simulation conditions. Our benchmarking also showed that false positives due to chain and fan-out relationships were best suppressed in EcohNet (Fig. 2i,j).

**Figure 3.**
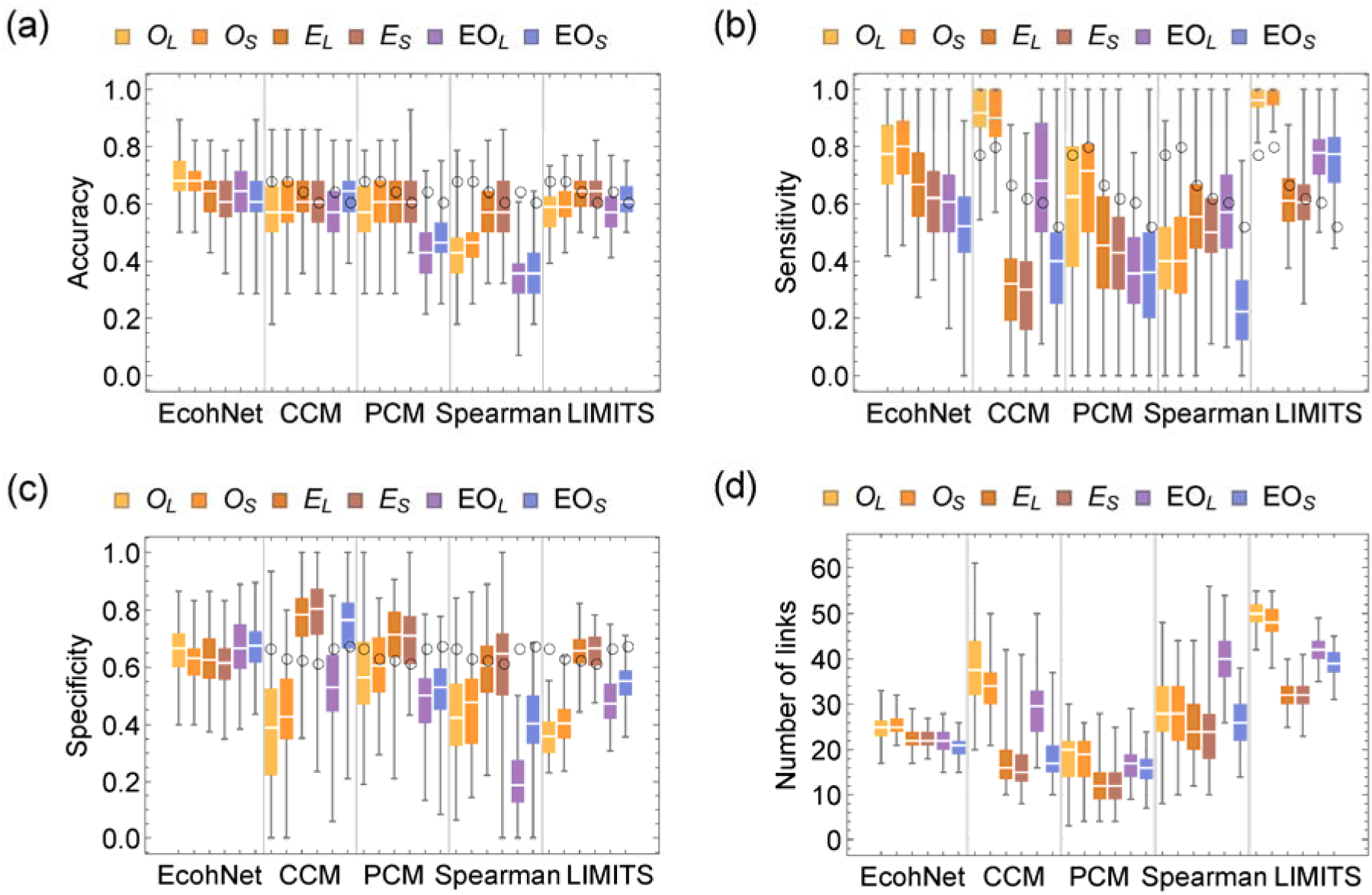
Values of accuracy (a), sensitivity (b), specificity (c) and number of inferred links (d) for the food web models. Since connectance was fixed at 0.33 when generating an interaction matrix, the number of links detected should ideally be a constant value (since the number of species was fixed at 8, average number of links was (8^2-8)*0.33=18). In the box plot, white lines indicate the median, box edges indicate the first and third quartile value, and whiskers indicate maximum and minimum values. Black circles indicate the median value of EcohNet.

The above conclusions did not change significantly based on different dataset sizes (SI Appendix Fig. S5,6), sampling intervals (SI Appendix Fig. S7-10), and the presence of unobserved species (SI Appendix Fig. S11) that we tested. While these results are based on food web models where the interactions are bidirectional, the superiority of EcohNet was shown also in a random interaction model where the interactions were unidirectional and the detectability of directional interactions were evaluated (SI Appendix Fig. S12; please see SI Appendix C for the evaluation of directional interactions in the food web models).

### Phytoplankton dynamics in a real ecosystem

We applied EcohNet to a time-series (Lake Kasumigaura Long-term Monitoring Dataset) to examine the top-down and bottom-up causal factors of phytoplankton community composition (Fig.4a). The causal network was naturally organized in a top-down structure with temperature at the top and temperature had the most numbers of interaction links across multiple trophic levels. EcohNet showed that each seven dominant phytoplankton groups were determined by different factors and those networks were complex. Four out of seven phytoplankton groups (all three cyanobacteria and Thalassiosiraceae) were forced by NO_3_-N. We also detected the top-down control of rotifers and calanoids on not only diatoms but also cyanobacteria (Rotifers→Nitzschia, Rotifers→Oscillatoriales, Calanoida→Microcystis, Clanoida→Thalassiosiraceae, Cyclopoida→Aulacoseira), while large- and small-cladocerans did not influence dominant phytoplankton groups. In contrast, three bottom-up links from phytoplankton to zooplankton (Nitzschia→Rotifers, Oscillatoriales→Cyclopoida, Fragilaria→Small-cladocera) were also detected. Importantly, in addition to the effects of environmental variables and zooplankton, we identified three interactions among dominant phytoplankton groups (Nostocales →Oscillatoriales, Oscillatoriales→Aulacoseira, and Fragilaria→Microcystis).

**Figure 4.**
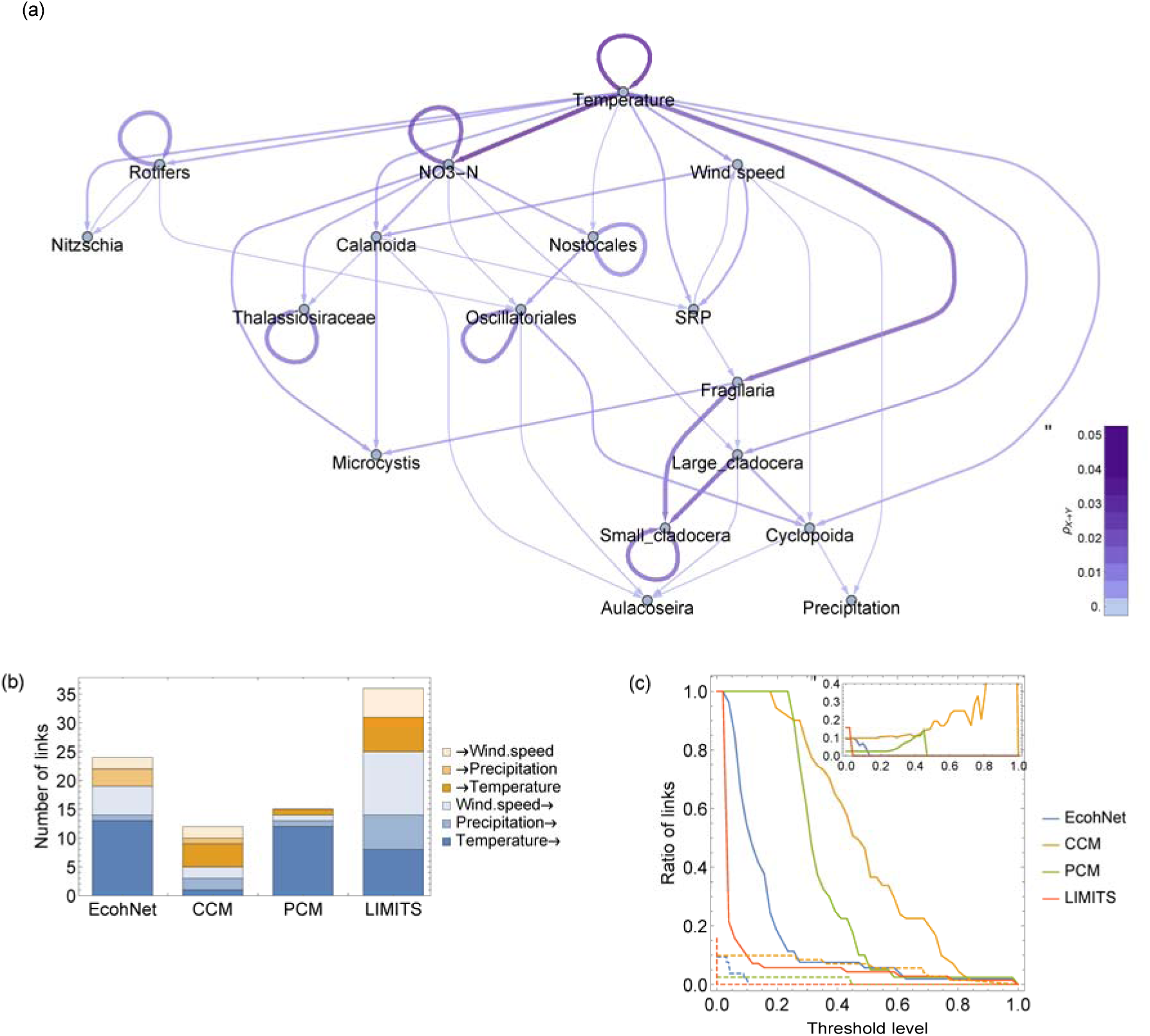
Analysis of Lake Kasumigaura Long-term Monitoring Dataset. (a) Causal network obtained by EcohNet. We removed causal links with a prediction skill of less than 0.002 (23 out of 70 links were removed). The magnitude of the prediction skill is indicated both by the thickness and color of arrows. Seven dominant phytoplankton groups were underlined. (b) We identified causal relationships including meteorological variables (Wind speed, Precipitation, Temperature) and examined how well unrealistic causal links (i.e., organisms/chemicals to meteorological variables) were avoided while the effect of meteorological variables were detected. (c) Ratio of causal links larger than the threshold level (x-axis) is shown for all links (solid lines) and unrealistic links (dashed lines). The inserted panel shows the ratio of the unrealistic links to all links. The faster the unrealistic/all ratio converge to zero, the smaller the unique prediction skill of unrealistic causal links, and the easier it is to remove them while retaining more reliable links.

An additional analysis on the direction of causation between meteorological and other components showed that the total number of unrealistic causalities was lowest in PCM, followed by EcohNet and CCM in this order (Fig. 4b). For the causal links with the meteorological variables as factors, EcohNet detected an equal number of links as PCM for temperature and precipitation and found more links for wind speed. Moreover, unrealistic causal links were likely to be detected as strong links in CCM and PCM, but appeared only as weak links in EcohNet (Fig. 4c).

## Discussion

We developed and evaluated the effectiveness of a method, EcohNet, that utilizes ensemble predictions of online echo state networks. In short, our method encompasses the scope of Granger causality [3] and nonlinear time series analysis [35], and implements a framework for decomposing relationships among variables in terms of predictability [23]. Although ESNs have been applied to causal analysis [36-38], we developed for the first time a reliable method of causal network inference by integrating adaptive online ESNs [20] into an ensemble machine learning framework. In addition to its applicability to nonlinear dynamics, our approach is not affected by the reliability of nonlinear prediction methods to the dynamics of a given system [39], which differs from previous approaches [5, 26, 28, 34, 40, 41]. As we have confirmed (Fig. 2e-g), performance of CCM and PCM, which relies on a specific nonlinear forecasting method (state space reconstruction and simplex-projection) was sensitive to the differences in dynamic complexity [42, 43], although it performed better than the Spearman rank correlation coefficient. Another advantage of our method is its capability to decompose relationships among variables; we confirmed that false positives due to chain and fan-out relationships were best suppressed in EcohNet followed by PCM (Fig. 2e,f). This result is plausible because PCM also address both relationships [28].

EcohNet has two prominent features. First, it was robust in ability to detect links and has relatively stable sensitivity and specificity (Fig. 3). This robustness is important because it means that the error rate of an inferred network is consistently controlled by the regularization scheme irrespective of the dynamics. For example, comparison of the connectivity of ecological networks from several sites would be difficult if the method applied was sensitive to the dynamical complexity [16]. Second, we confirmed that EcohNet can identify causation between meteorological and other components while filtering out the unrealistic direction of causalities (Fig. 4b,c) [18]. The unrealistic causal links were likely to be detected only as weak links in EcohNet. Because of observational noise and other factors, it is hard to completely avoid detecting unrealistic causalities. Therefore, it is a desirable feature that the importance of unrealistic causality is underestimated. Among the rest of the methods, PCM had stable sensitivity and specificity and suppressed the detection of unrealistic links. On the other hand, its overall performance (Fig. 2g,h) was no better than CCM. One reason would be the absence of steps for evaluating the convergence of a prediction skill in PCM whereas in CCM this helps to discount the predictability of the target variable itself. In EcohNet, it is considered as *ρ*_*X*→*X*_.

EcohNet revealed that temperature determined nutrient dynamics, phytoplankton, and zooplankton communities in Lake Kasumigaura. This result is similar to Tanentzap et al. [44] that demonstrates strong direct and indirect impacts of temperature on phytoplankton and zooplankton. Quantifying the causal effects of temperature is an important advantage of EcohNet. CCM failed to distinguish causal relationships from seasonality-driven synchronization, leading to misidentification of causality [45]. Likewise, previous work on Lake Kasumigaura did not detect the causal effects of temperature on phytoplankton and zooplankton [25]. Our results suggest that temperature may determine the whole food web structure of Lake Kasumigaura.

Clearly, each seven dominant phytoplankton groups were determined by different factors (Fig. 4). Some studies reported that phytoplankton species or genera have unique physiological and ecological features and thus respond differently to environmental factors and grazing [46, 47]. These differences can result in dynamic changes in phytoplankton community composition, since it is reported that the dominant phytoplankton group changed temporally in Lake Kasumigaura [48-50]. NO_3_-N determined four out of seven phytoplankton groups. This is consistent with previous work on Lake Kasumigaura showing nitrogen limits phytoplankton primary production [25]. Not only diatoms but also cyanobacteria were influenced by rotifers, calanoids, and cyclopods. Although our analysis did not identify whether these are direct predation or indirect effects, it is known that some copepods can ingest and shorten the filament size of cyanobacteria and rotifers can graze or utilize decomposed cyanobacteria [51, 52]. Our results suggest that these zooplankton groups might have an important role in the food web of shallow hypereutrophic lakes.

We identified significant interactions among dominant phytoplankton groups. It has been reported that phytoplankton groups may have compensatory responses to environmental factors [53]. One possible mechanism for the interaction between Anabaena and Oscillatoriales could be nitrogen availability, because N_2_-fixing cyanobacteria, including Nostocales, compete with non-N_2_-fixing bacteria, including Oscillatoriales [54]. Another mechanism could be light availability. Mixing regimes determined by climatic factors like heat exchange and wind action affect competition for light between phytoplankton species (e.g., buoyant cyanobacteria and sinking diatoms), and shading by cyanobacteria blooms also influences other phytoplankton [55]. As pointed out in Freeman et al. [47], previous studies have not incorporated these interactions in models for identifying species-specific factors. Explicitly including not only environmental factors and grazing impacts but also community interactions in complex systems is another advantage of EcohNet.

As well as predicting the flow of causation, EcohNet enables cyanobacteria bloom forecasts for a lake (SI Appendix Fig. S13, 14). The application of EcohNet to long-term monitoring data can be a useful tool for lake management. Harmful cyanobacteria blooms are a serious threat to ecological integrity and ecosystem services, despite management efforts, and climate change is predicted to promote the occurrence and severity of cyanobacterial blooms [56]. As show here, the system is complex and therefore, identifying the drivers causing cyanobacteria blooms and forecasting these blooms using EcohNet are critically important for water quality management [29].

There are two developmental directions that we did not fully address in this paper. First, in this paper we did not consider the possibility of synergistic effects of two or more variables that may contribute more to the prediction than simple addition of their effect. Of course, we know this can occur [57] but we believe that a further extension of our methodology, e.g., evaluating the combinatorial effect of removing multiple variables, could address this issue. Second, we did not consider data-dependent parameter fitting of ESN-RLS. It may be true that optimal settings of parameters would maximize the performance of EcohNet’s network inference. Guiding principles for such a setup would be to use autocorrelation to determine the forgetting coefficient *λ* [58], and to determine parameters to maximize the predictability of the time series [59]. However, we have demonstrated the benefits of EcohNet well enough without these further optimizations.

The development of frameworks to facilitate time-series-based causal analysis would advance the field of ecology in the coming decades, where advanced techniques for ecosystem monitoring [60-64] will lead to unprecedented increases in dataset size [7, 65, 66]. Such analysis could promote studies to estimate the effects of global environmental change on biological communities [27], prevent regime shifts that lead to catastrophic impacts on biodiversity [67], and support efficient management of biological resources [68]. Further development by applying EcohNet to time-series of high-frequency data, such as sensor data, could lead to real-time, near-term forecasting to prepare for or preempt future impairment of ecological functions and services. It is expected that with integration to active hypothesis testing, these studies will aid the development of predictive and manipulative ecology in the 21st century.

## Materials and Methods

H2>Echo state network with a recursive least squares algorithm (ESN-RLS)

An echo state network (ESN) [19, 21] implements a type of reservoir computing that uses a recurrent neural network (RNN) as a dynamical reservoir (an internal structure which is made up of individual, non-linear units, and can store information). ESNs can skillfully reconstruct and predict time series from different nonlinear dynamical systems [69-75]. It has been demonstrated that ESNs can track the temporal evolution of time series better than backpropagation-based artificial neural networks [71].

In an ESN, an input signal induces a nonlinear response to the dynamical reservoir RNN, and the reservoir states are converted to an output signal by linear weights (Fig. 1C). Here, for a multivariate time series *X*= {*x*_1_, *x*_2_,…, *x*_*n*_}, we assume that *X*^Ω^(*t*) = {*x*_*λt*_}_*λ*∈Ω_is the input at time *t* specified by a set of indices Ω⊂{1,2, …, n}, and *x*_*i*_= (*x*_*i*1_, *x*_*i*2_,…) ∈ *X* is a target variable to which an ESN is trained to output its one-step-ahead prediction. Following the typical implementation, we defined an ESN by three matrices and one function. The matrices are: 1) the input weight matrix, 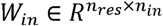; 2) the reservoir weight matrix, 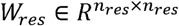 and 3) the output vector, 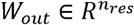, where *n*_*in*_, *n*_*res*_ is the number of nodes in the input layer (number of elements in Ω), dynamical reservoir, respectively. The state update equations for an ESN are as follows:

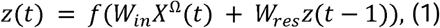

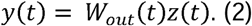

Here, *z*(*t*) is a vector of neural states in the dynamical reservoir, *y*(*t*) is the output of an ESN as the prediction of *x*_*i*_(*t* + 1), and *f*(*z*) = *tanh z* is a neural activation function. The neural connections of an ESN are randomly generated, except for the output weights *W*_*out*_. Here, *W*_*out*_is sequentially updated by the RLS (recursive least squares) method. This defines an online implementation of ESN, namely, ESN-RLS, which is robust to the non-stationary dynamics and easily applicable to predictive purposes [20].

RLS is widely used in linear signal processing and has a desirable feature of fast convergence [76]. It incorporates the error history of a system into the calculation of the present error compensation. The recursive updating rule of RLS for *W*_*out*_is as follows:

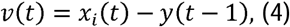

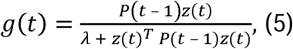

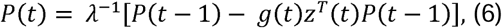

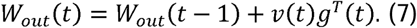

Here, *λ* is the forgetting factor, which determines the effect of past errors on the update, *v*(*t*) ∈*R* is the output error at time *t* when applying the weight matrix before the update, 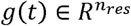 is the gain vector, and 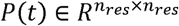 is the inverse of the self-correlation matrix of *z*(*k*) (*k* = 1,2, …, *t*) weighted by the forgetting factor. This update can be repeated multiple times at each time step, and we define the number of iterations as *τ*. We set the initial conditions of the update rule as,

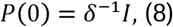

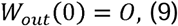

where *I* is a *n*_*res*_× *n*_*res*_identity matrix, *O* is a zero-vector of length *n*_*res*_and *δ* is the regularization factor. We also set the initial reservoir state as *z*(0) = *O*.

This algorithm minimizes the following cost function, which is a weighted sum of the output errors at time *k* = (1,2,…, *t*) when applying the output weight matrix *W*_*out*_at time *t* with a regularization term:

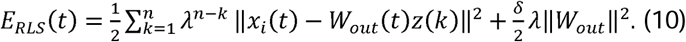

Here, ||·|| represents the L_2_ norm.

### Parameterization of ESN-RLS

The parameter values were set according to the standard recommendation for parameterization of the ESN and RLS [20, 21 76-78] (Table 1, SI Appendix B). The elements of *W*_*in*_are randomly drawn from a uniform distribution between −0.1 and 0.1. We scaled the spectral radius of *W*_*res*_by *ρ*_0_after it is generated by randomly drawing its elements from a uniform distribution between −1 and 1 (with probability 0.1, and otherwise zero). The number of nodes in the dynamic reservoir *n*_*res*_was 32, which is smaller than that of a typical ESN. This is intended to reduce computation time. Also, this number is related to the memory capacity of the reservoir [20], and the number of nodes must be scaled to the time series length. In this sense, a large number of nodes was not needed in this study. In our benchmark using aquatic microcosm data, we examined the stability of the results for four representative parameters (*ρ*_0_, *λ, τ*., *δ*) that may affect the performance of ESNs (SI Appendix Fig. S16).

**Table 1.** Parameters of ESN-RLS. For the parameters marked by *, we tested the impact on EcohNet’s performance with different values. The tested values are τ ∈ {2,4,6,8,10}, *λ* ∈ {0.8,0.9,0.95,0.98,0.995}, *δ* ∈ {0.0001,0.001,0.01,0.1}, *ρ*_0_ ∈ {0.7,0.8,0.9,0.95,0.98}.

| Description | Value |
| --- | --- |
| Connectance of $W_{in}$ | 1 |
| Range of values in $W_{in}$ | [-0.1,0.1] |
| Number of nodes in $W_{res}$ ($n_{res}$) | 32 |
| Connectance of $W_{res}$ | 0.1 |
| Range of values in $W_{res}$ | [-1,1] |
| Spectral radius of $W_{res}$ ($\rho_0$) | 0.95* |
| Forgetting factor ($\lambda$) | 0.95* |
| Number of iterations of RLS updates ($\tau$) | 8* |
| Regularization factor ($\delta$) | 0.001* |

### Progressive selection of input variables

For a target variable *x*_*i*_(corresponds to X in Figure 1A), we used a progressive selection to obtain the set of variables 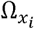 that minimizes the prediction skill for *x*_*i*_({X, Y} in Fig. 1B). For this purpose, we first define the prediction skill in this paper as:

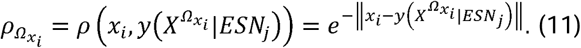

Here, *x*_*i*_is the time series of target variable (blue lines in Fig.1C) and 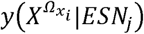 represents the prediction of *x*_*i*_given a set of variables 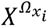 and an echo state network indexed by *j*(yellow lines in Fig.1C). Practically, we calculated the prediction skill after truncating the first 20% of the time series. It is intended to remove the effect of initial condition (*z*(0), *P*(0) and *W*_*out*_(0)).

To prevent the results from being dependent on a particular network, the evaluation of prediction skill is done by an ensemble of *N* ESNs. This allows us to account for the estimation error in prediction skill and ensures more reliable variable selection. Moreover, to avoid overfitting, the input is replaced by a zero vector with a fixed probability at each time. This means that we eventually use the following equation to update the reservoir state instead of eq. (1):

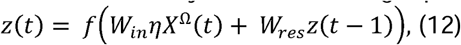

where *η* is a random number that is 1 with probability 0.5 and 0 otherwise.

The progressive selection of variables proceeds as follows. First, the set of indices of input variables to predict variable *x*_*i*_is initialized to *I*_*active*_= {*i*} to account for the predictability inherent in the target variable itself (first step in Fig.1B). Correspondingly, the set of indices of the rest of the variables is initialized to *I*_*inactive*_= {*j*}_*j*≠*i*_. Then, *N* ESNs are generated to perform predictions of *x*_*i*_ by setting 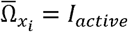 = *I*_*active*_ (Fig.1C). This returns a distribution of prediction skills *FT*= 0(*T* indicates the step of variable selection). To select the next variable to be added, for each index 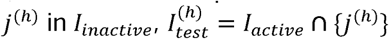 are generated, where the suffix *h* indicates that j^*(h*)^is the *h*th element of *I*_*inactive*_, and then, for each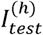, *N* ESNs are generated to obtain predictions of *x*_*i*_ by setting 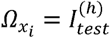 (2nd step in Fig.1B). Among the distribution of the prediction skills for *h*S, onethat has the highest median value (*h*= *h*^*^) is set as *F*_*T*+1_(second step in Fig.1B). The criterion used to accept *F*_*T*+1_as 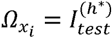 is that the median of *F*_*T*+1_is larger than *q* × 100% of values in*F*_*T*_, formally written as, #{*ρ* ∈ *F*_T_|*ρ* > *median*(*F*_T +1_)}*IN* < *q*, where #{*ρ* ∈ *F*|*criteria*} is the number of *ρ* in *F* that satisfies the *criteria*. If the criterion is satisfied, we replace *F*_T_by *F*_T+1_, *I*_*active*_by 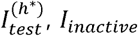 by *I*_*inactive*_ \ {*j*(*h*\*)} (removing *j*^(*h*\*)^from *I*_*inactive*_), increment *T* by 1, and proceed tothe next step only if *I*^*inactive*^≠ {}, otherwise the procedure is stopped (3rd step in Fig.1B) with returning 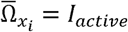 as the set of the indices of optimal variables that maximizes the predictionof *x*_*i*_.

The number of ESNs generated for each evaluation, *N*, is related to the stability of the results. If *N* is small, the variability in results from trial to trial may be large, especially for weak causality. In this paper, we set *N* = 10000 for real-world data to obtain stable results, and *N* = 5000 for simulation data sets to reduce the time required for evaluation. The threshold value for accepting a new variable, *q*, was set to 0.48 throughout this paper. The closer the number is to 0.5, the more likely it is that weak links will remain. Therefore, instead of increasing the sensitivity (true positive rate), there is a possibility of decreasing the specificity (increasing the false positive rate). In this paper, the value is set close to 0.5, taking into account the evaluation by ROC-AUC.

### Calculation of unique prediction skill

Unique prediction skill quantifies the unique contribution of a single variable to the overall prediction skill (Fig. 1D). For each 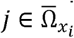, the unique prediction skill is defined as,

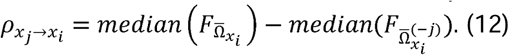

Here, *F*_Ω_is the distribution of prediction skill for Ω and,

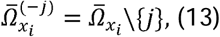

i.e., 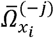 is obtained by removing *j* from 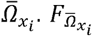 and 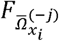 is again calculated by *N* ESNs. 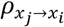 represents how *x*_*j*_uniquely contributes to the prediction of *x*_*j*_(illustrated as the difference of the positions of two distributions in Fig.1D). We identify 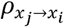 as the indicator of causal influence from *x*_*j*_to *x*_*i*_. In this step, 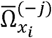 such that 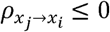 is removed from 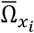, and we set 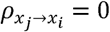 as well as other variables that does not included in 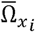.

### Dataset

We benchmarked our method using both experimental and simulation data (SI Appendix D) and then applied it to a long-term monitoring dataset from a lake ecosystem. One of the benchmarking datasets was from a real ecological system, which was obtained by long-term observation of a mesocosm [33]. The other datasets were obtained by simulations of two ecological models, namely food-web and random interaction model (all simulation conditions are listed in SI Appendix Table S1), and allowed us to examine how dynamic complexity of a time series affect identification of causal relationships, and how the effects of two types of interaction topologies (chain and fan-out relationships) can cause false positives.

### Lake Kasumigaura Long-term Monitoring Dataset

We applied the EcohNet to the long-term monitoring data from Lake Kasumigaura, a hypereutrophic lake. Lake Kasumigaura is the second largest lake in Japan (167.7 km^2^) and is shallow (mean depth: approximately 4 m, maximum depth: 7.4 m). The National Institute for Environmental Studies has been conducting monthly monitoring in Lake Kasumigaura since 1976 and publishing the long-term data on the Lake Kasumigaura Database. In this lake, cyanobacteria blooms occur and disappear repeatedly, and the dominant cyanobacteria group changes.

Phytoplankton communities can interact with many components, such as temperature, nutrients, and zooplankton, and are complex and dynamic. Using EcohNet, we quantified the causal relationships among environmental variables, seven dominant phytoplankton groups (three cyanobacteria (Microcystis, Nostocales, Oscillatoriales) four diatoms (Thalassiosiraceae, Aulacoseira, Fragilaria, Nitzschia)), and zooplankton, and examined the interactions among these phytoplankton groups.

We analyzed the monitoring data at the center of Takahamairi Bay (Station 3), which is shallow (ca. 3.2 m) and the most eutrophic site with cyanobacteria blooms. Phytoplankton biovolume data was obtained from Takamura and Nakagawa [79]. For zooplankton, we used the abundance data of five functional groups: large cladocerans (>1.0 mm), small cladocerans (<1.0 mm), rotifers, adult calanoids, and adult cyclopoids [25]. Zooplankton data were obtained from Takamura et al. [80]. Our key environmental variables were surface water temperature, soluble reactive phosphorus (PO_4_-P), and nitrate nitrogen (NO_3_-N) from the Lake Kasumigaura Database (https://db.cger.nies.go.jp/gem/moni-e/inter/GEMS/database/kasumi/index.html). We also included 30-day moving average of wind speed and precipitation, which were collected from the Tsukuba-Tateno Meteorological Station of the Japan Meteorological Agency (https://www.jma.go.jp/jma/menu/menureport.html). Since CCM, PCM and LIMITS requires taking consecutive time lags of observed variables, we analyzed data from April 1996 to March 2019 (276 months). There were no missing data for any variables during this time interval. All time series were square-root-transformed and normalized to have a mean 0 and variance of 1 to adjust for rapid increases in some phytoplankton species.

### Evaluation

We evaluated the performance of EcohNet and conventional methods (SI Appendix A) in detecting interactions using the evaluation criteria of a binary classification task, namely, ROC-AUC (the area under the curve of a receiver-operator characteristic curve), F1 score, accuracy, sensitivity and specificity (SI Appendix Fig. S3). Since accuracy is affected by the degree of connctance, we considered ROC-AUC and F1-score as criteria for overall performance. We specifically considered prediction skill (EcohNet, CCM and PCM), correlation coefficient (Spearman rank correlation) and interaction coefficient (LIMITS) as a classifier to identify interactions. We adjusted the sparsity of the Spearman rank correlation matrix based on the p-value, i.e., we made the correlation matrix sparse by replacing elements with *P* > 0.01 with 0. In the same manner, we set the threshold p-value of CCM as 0.05 (SI Appendix A2). For PCM, following the author implementation [28], we used the value of prediction skill instead of p-value and set the threshold as 0.2. For LIMITS, as in EcohNet, the sparsity of an interaction matrix depends on the forward stepwise algorithm. Here, the threshold value for the improvement of prediction error (SI Appendix A4) was set as 0, i.e., new variable is added if it at least improves the prediction.

In predator-prey relationships species affect each other directly, whereas in a causal relationship, a relationship between species may only be detected in one direction due to a difference in the time-dependence of each effect [5, 24, 25]. Therefore, in this paper, when evaluating EcohNet, CCM and PCM for the long-term mesocosm experiment and the food web models using the evaluation criteria of the binary classification task, we consider the ability to detect interacting pairs (pairs where at least one has a direct influence on the other). Specifically, we symmetrized the matrices of the prediction skill of EcohNet, CCM and PCM so that *a*_*ij*_, *a*_*ji*_← *max*(|*a*_*ij*_|, |*a*_*ij*_|) before evaluation, and evaluated the performance of prediction skill as a classifier in detecting elements for which *I* < *j* and whose values are non-zero in the actual interaction matrix. The matrix of Spearman rank correlation, which is originally symmetric, was evaluated in the same manner. For the random interaction model, we considered the ability of EcohNet, CCM and PCM to detect direct influences. Specifically, we tested the presence of influence from one species to another (corresponding to a non-zero *a*_*ij*_in the interaction matrix) when there are causal links from the former to the later. For LIMITS, we evaluated the performance of detecting non-zero components of the interaction matrix directly in all cases according to its original definition.

## Data availability

We used Mathematica 12.3 and 13.1 for our analysis. The computer codes used for the analysis can be downloaded from: https://github.com/kecosz/EcohNet (DOI: 10.5281/zenodo.5492161)

## Acknowledgments

We especially thank Shinji Nakaoka for valuable discussions, and Megumi Nakagawa for counting phytoplankton and providing helpful comments. We also thank all members of the Lake Kasumigaura long-term monitoring group of the National Institute for Environmental Studies. This work was supported by funding from the Management Expenses Grant for RIKEN BioResource Research Center, MEXT, and the Japan Society for the Promotion of Science (JSPS) KAKENHI JP20K06820 and JP20H03010 (to K.S.).

