## Supplementary material for "Decomposing predictability to identify dominant causal drivers in complex ecosystems": SI Appendix

SI Appendix for

SI Appendix Figures

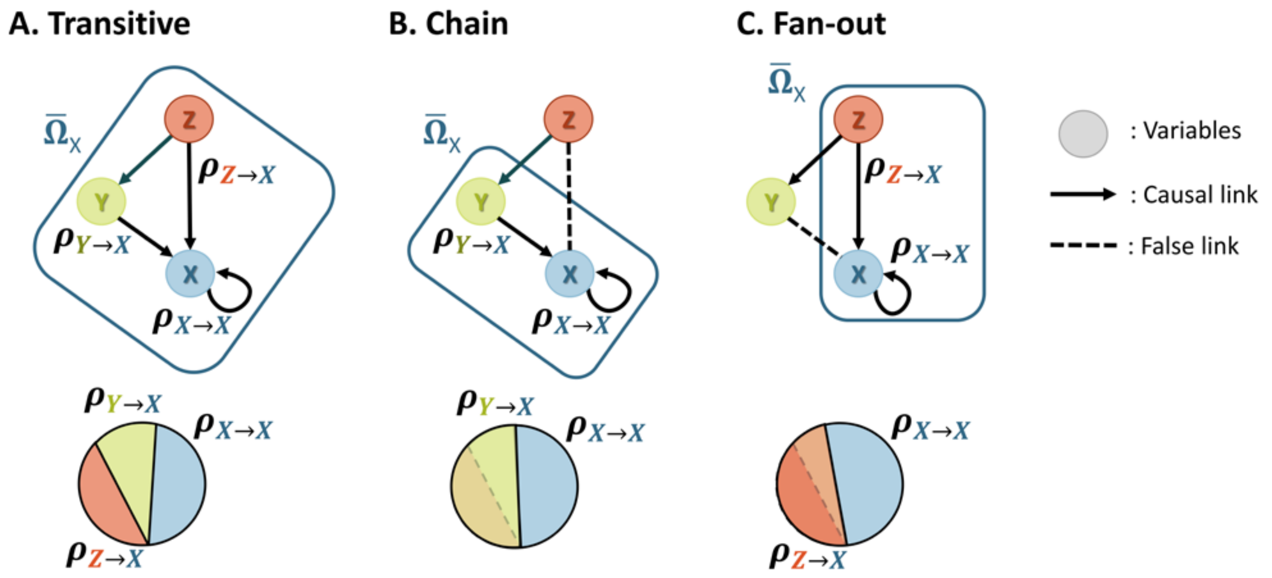

Figure S1. Schematic description of unique prediction skill in transitive (A), chain (B) and fan-out (C) relationships. In (A), both Y and Z have unique predictive skills for X because there is direct relationship between X and Y. However, in (B) and (C), only either Y or Z, respectively, is expected to have non-zero value in addition to X, if all else is equal. In (B), because of the absence of direct relationship between X and Z, the contribution of Z on X is always mediated by Y and thus only $\rho_{Y \to X}$ and $\rho_{X \to X}$ is expected to have non-zero value. In (C), because X and Y share influence from Z, the direct relationship between X and Z covers contribution of Y on X and thus only $\rho_{X \to X}$ and $\rho_{Z \to X}$ is expected to have non-zero value.

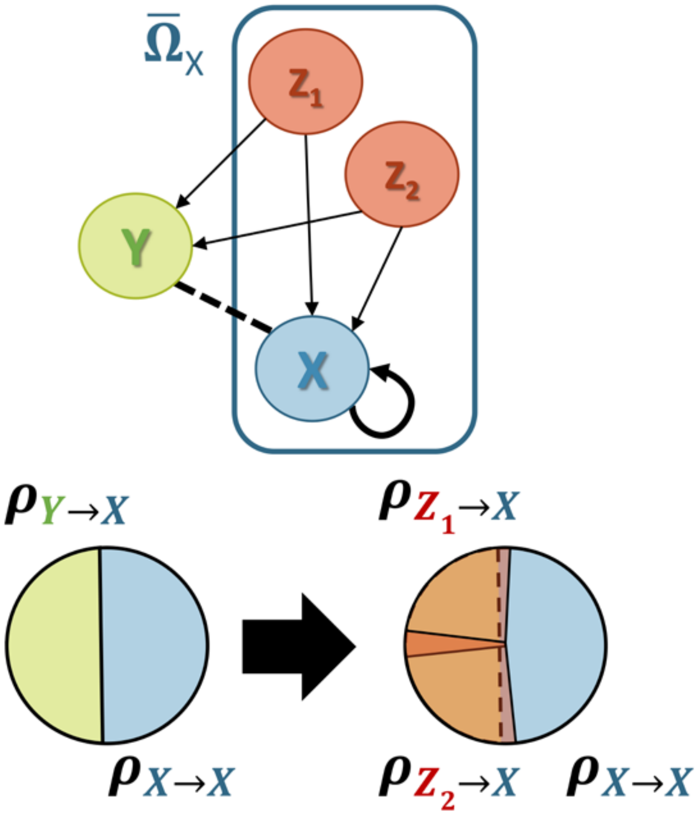

Figure S2. Both $Z_1$ and $Z_2$ have a fan-out relationship with X and Y. A simple forward stepwise algorithm would select Y first followed by $Z_1$ and $Z_2$, because Y has the larger individual contribution than either $Z_1$ or $Z_2$. However, the evaluation of unique prediction skill can remove Y from $\bar{\Omega}$ since contribution of Y is included in the union of the contribution of $Z_1$ and $Z_2$.

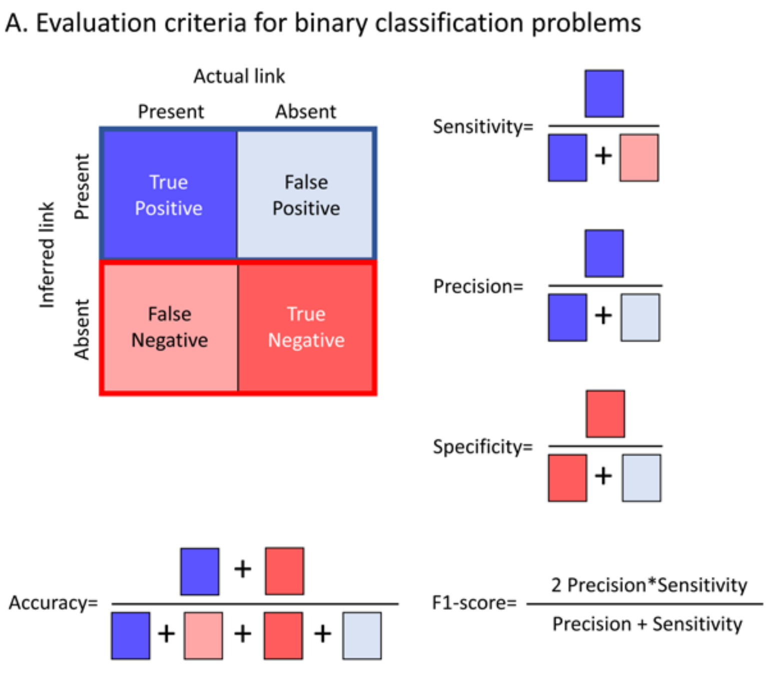

Figure S3. Evaluation criteria for binary classification problems and their relationship to ROC curves. (A) evaluation criteria for binary classification problems. True positive (TP) represents a result of inference that correctly indicates the presence of links, true negative (TN) represents a result of inference that correctly indicates the absence of links, false positive (FP) represents a result of inference which incorrectly indicates presence of links, and false negative (FN) is a result of inference which incorrectly indicates the absence of links. (B) Explanation of ROC curve. Here, a circle corresponds to an interaction between species. Suppose we set a threshold value for a classifier and determine that links between species above the threshold value are present, and links below the threshold value are absent. By matching this with the actual presence/absence status of interaction links, we can calculate sensitivity and specificity and plot a single point on the 2D plane. The ROC curve is obtained by repeating the calculation with changing threshold values, and the ROC-AUC is the area underneath the curve.

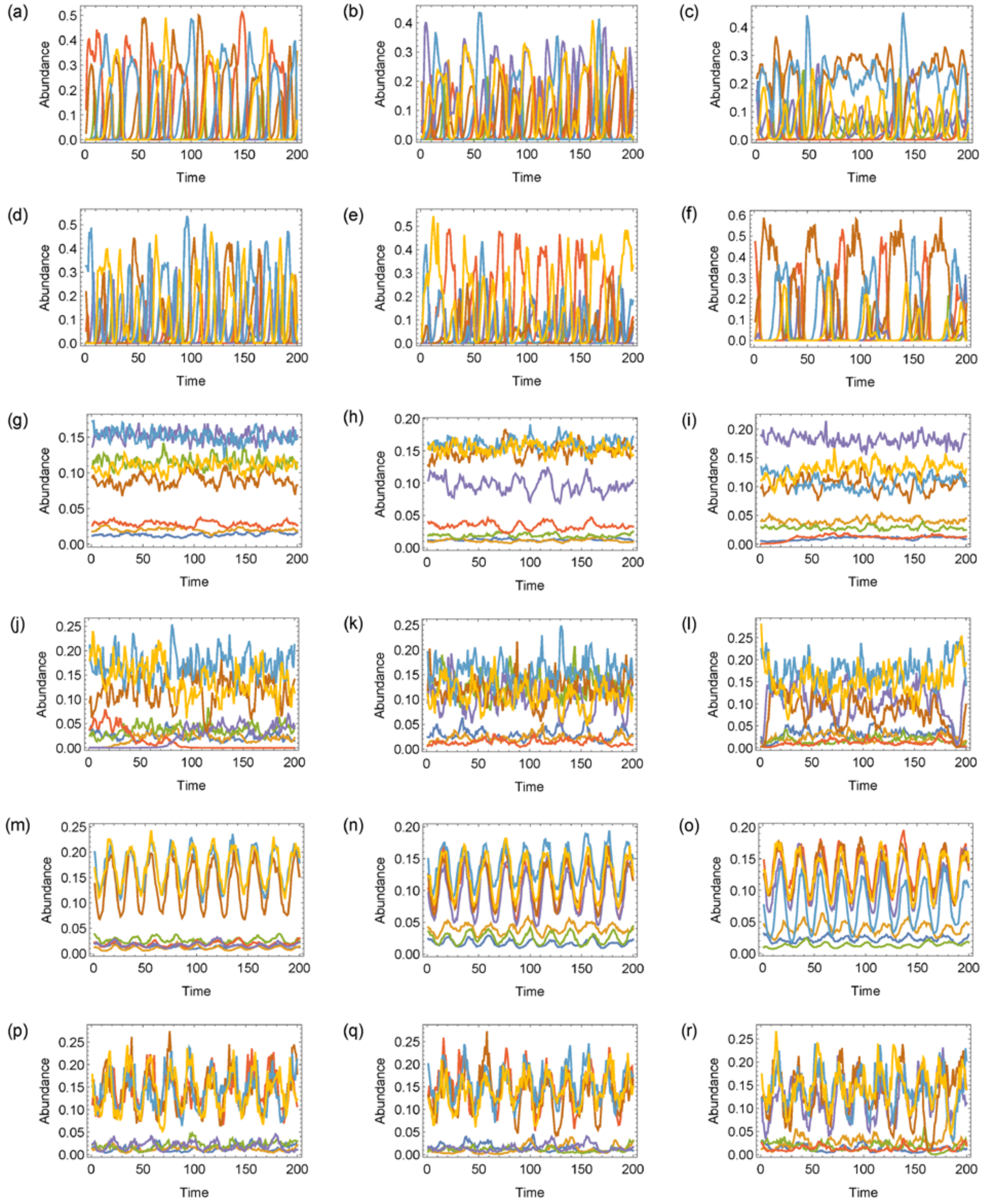

Figure S4. Representative time series of six different models in simulation dataset. (a-c) $O_S$, (d-f) $O_L$, (g-i) $E_S$, (j-l) $E_L$, (m-o) $EO_S$, and (p-r) $EO_L$.

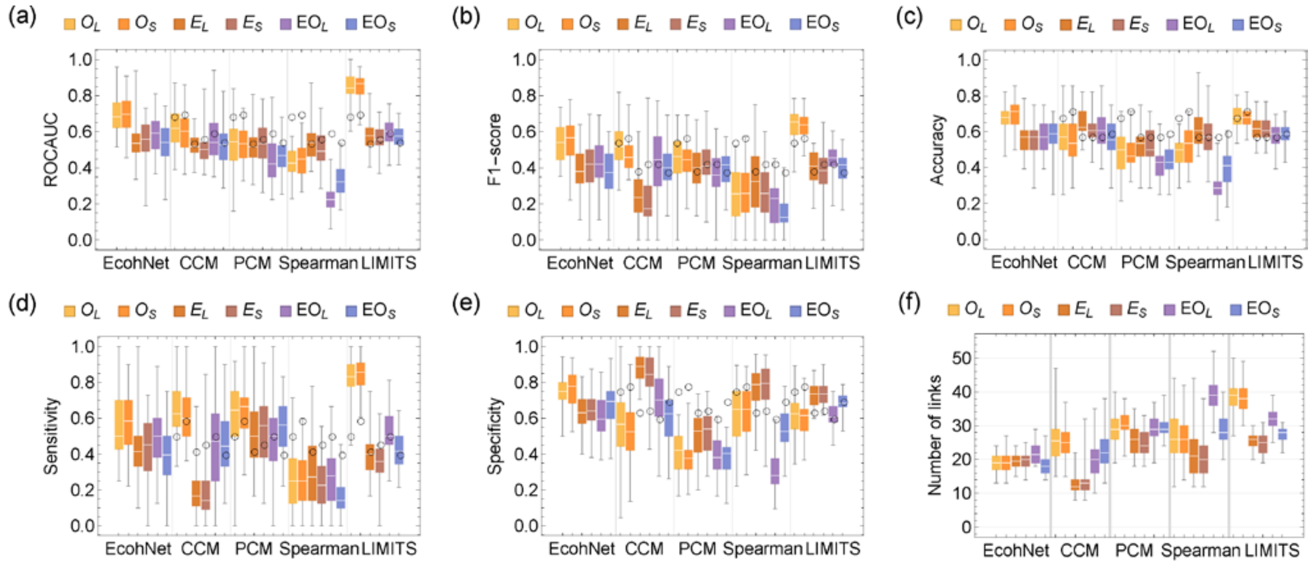

Figure S5. The value of ROC-AUC (a), F1-score (b), accuracy (c), sensitivity (d), specificity (e), and the number of inferred links (f) for the food web models (2,000 steps sampled with an interval of 40 steps). In the box plot, white lines indicate the median, box edges indicate the first and third quartile value, and whiskers indicate maximum and minimum values. Black circles indicate the median value of EcohNet.

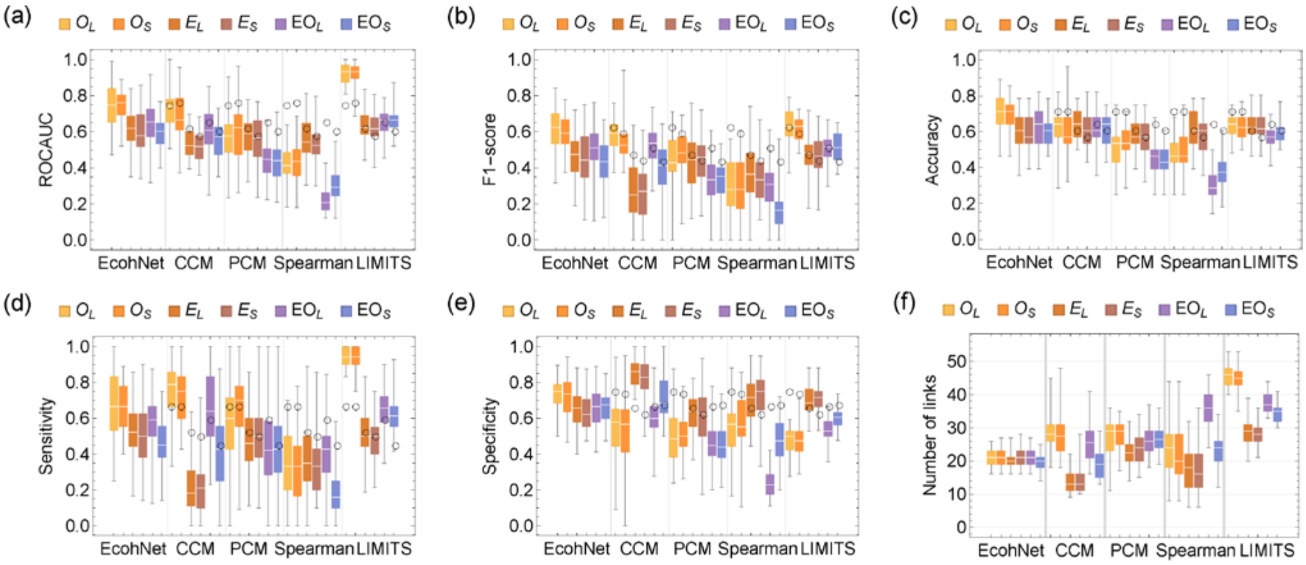

Figure S6. The value of ROC-AUC (a), F1-score (b), accuracy (c), sensitivity (d), specificity (e), and the number of inferred links (f) for the food web models (4,000 steps sampled with an interval of 40 steps). In the box plot, white lines indicate the median, box edges indicate the first and third quartile value, and whiskers indicate maximum and minimum values. Black circles indicate the median value of EcohNet.

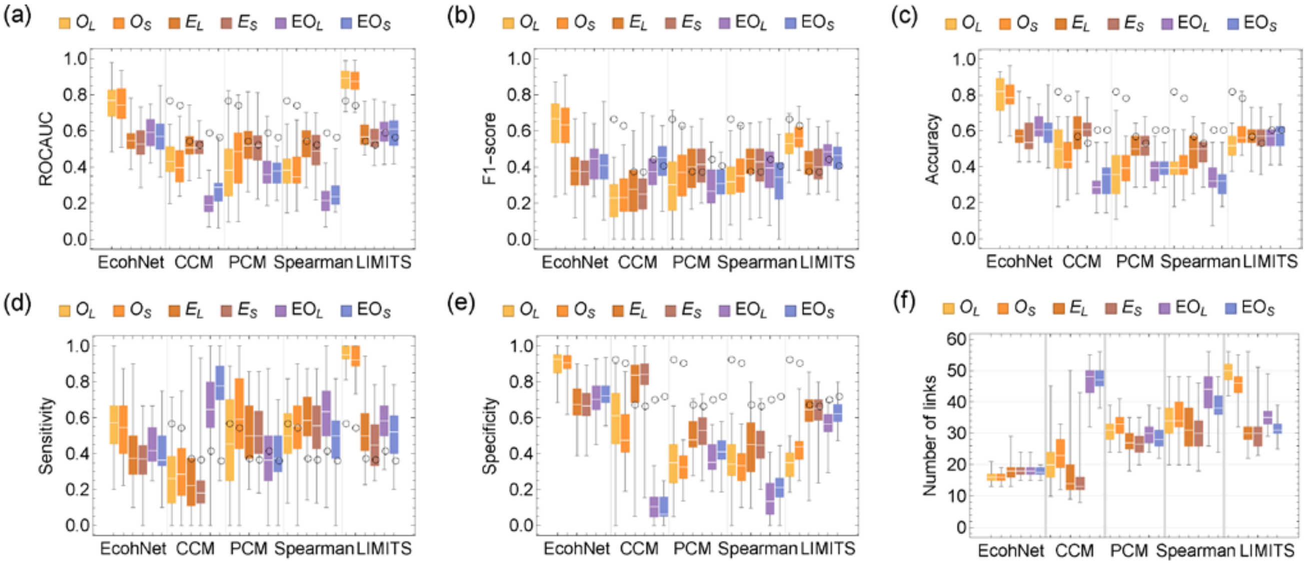

Figure S7. The value of ROC-AUC (a), F1-score (b), accuracy (c), sensitivity (d), specificity (e), and the number of inferred links (f) for the food web models (2,000 steps sampled with an interval of 10 steps). In the box plot, white lines indicate the median, box edges indicate the first and third quartile value, and whiskers indicate maximum and minimum values. Black circles indicate the median value of EcohNet.

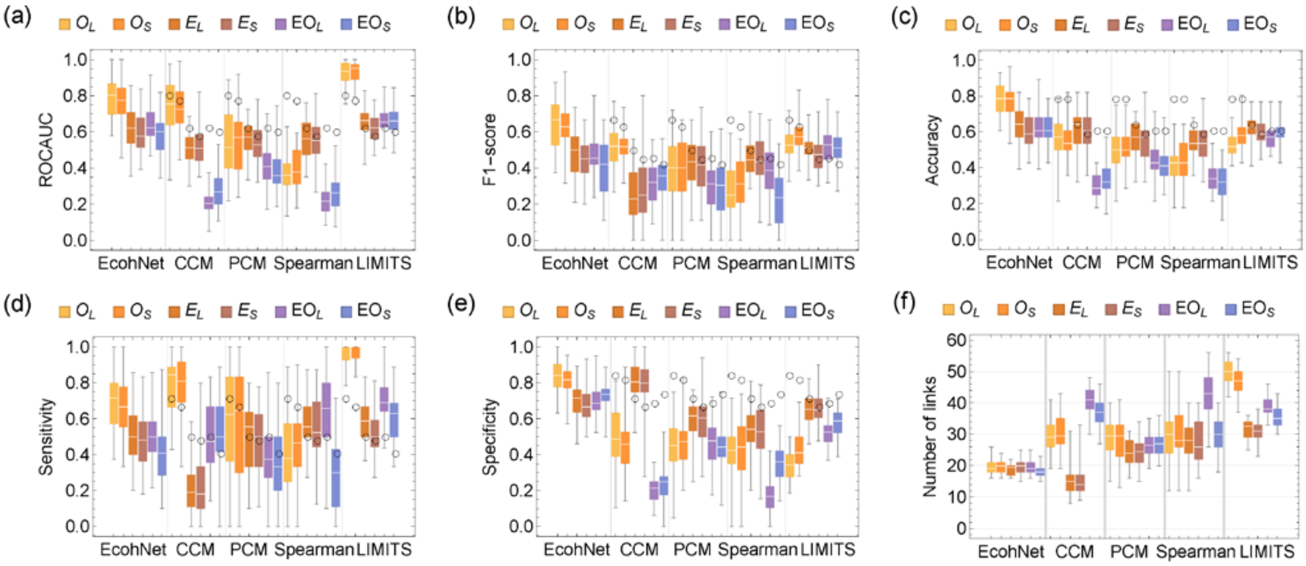

Figure S8. The value of ROC-AUC (a), F1-score (b), accuracy (c), sensitivity (d), specificity (e), and the number of inferred links (f) for the food web models (4,000 steps sampled with an interval of 20 steps). In the box plot, white lines indicate the median, box edges indicate the first and third quartile value, and whiskers indicate maximum and minimum values. Black circles indicate the median value of EcohNet.

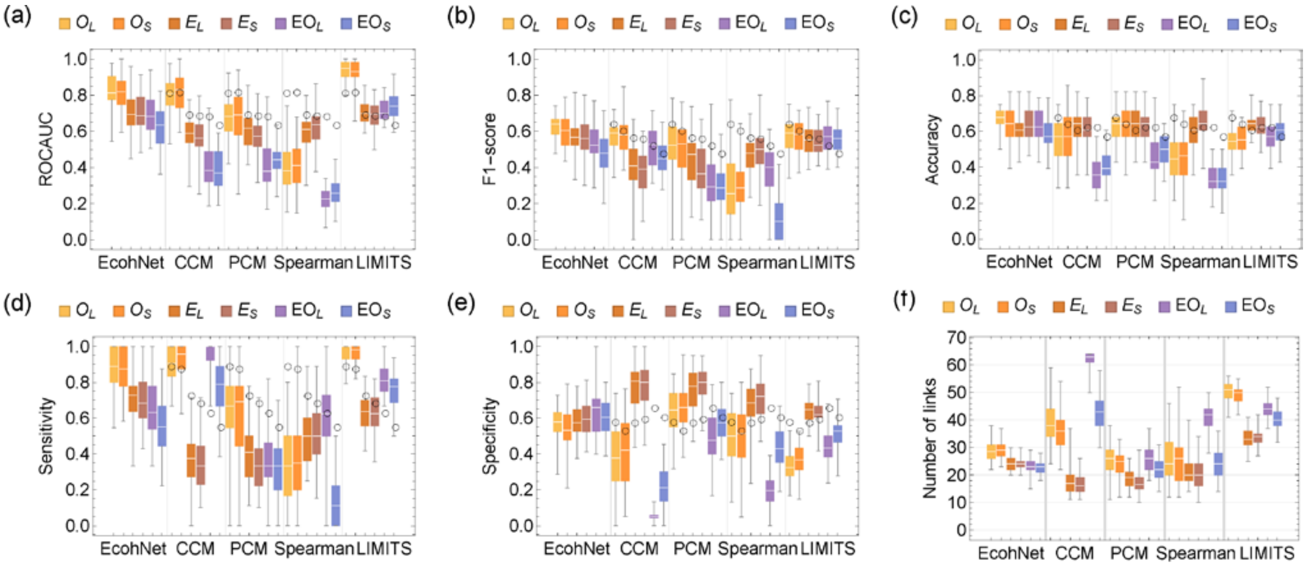

Figure S9. The value of ROC-AUC (a), F1-score (b), accuracy (c), sensitivity (d), specificity (e), and the number of inferred links (f) for the food web models (12,000 steps sampled with an interval of 60 steps). In the box plot, white lines indicate the median, box edges indicate the first and third quartile value, and whiskers indicate maximum and minimum values. Black circles indicate the median value of EcohNet.

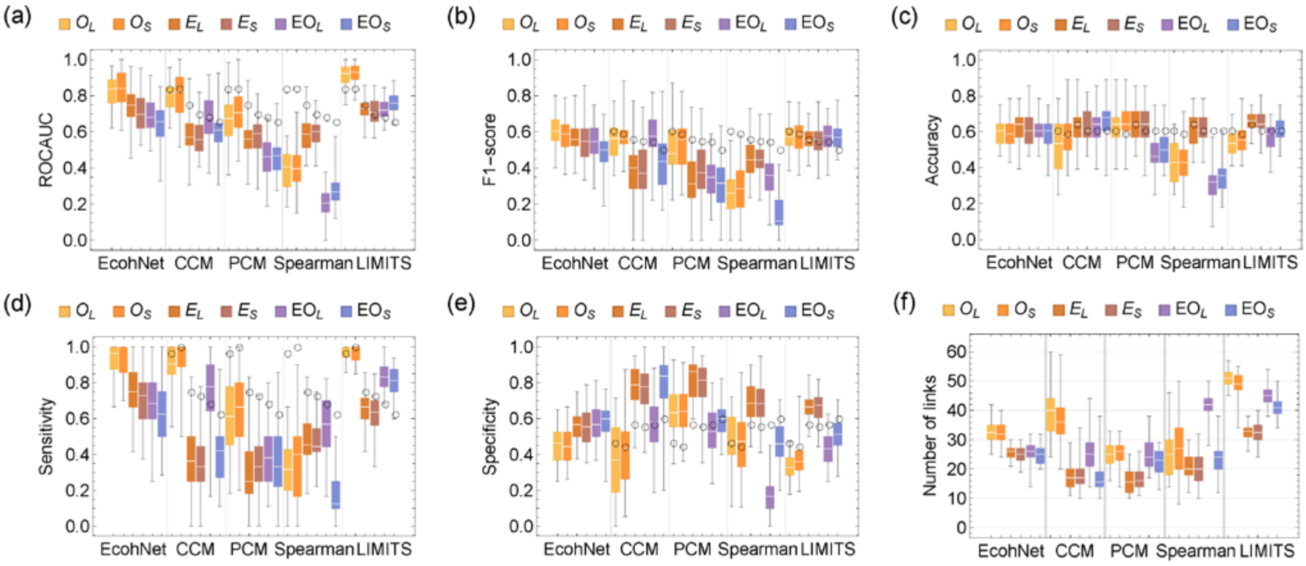

Figure S10. The value of ROC-AUC (a), F1-score (b), accuracy (c), sensitivity (d), specificity (e), and the number of inferred links (f) for the food web models (16,000 steps sampled with an interval of 80 steps). In the box plot, white lines indicate the median, box edges indicate the first and third quartile value, and whiskers indicate maximum and minimum values. Black circles indicate the median value of EcohNet.

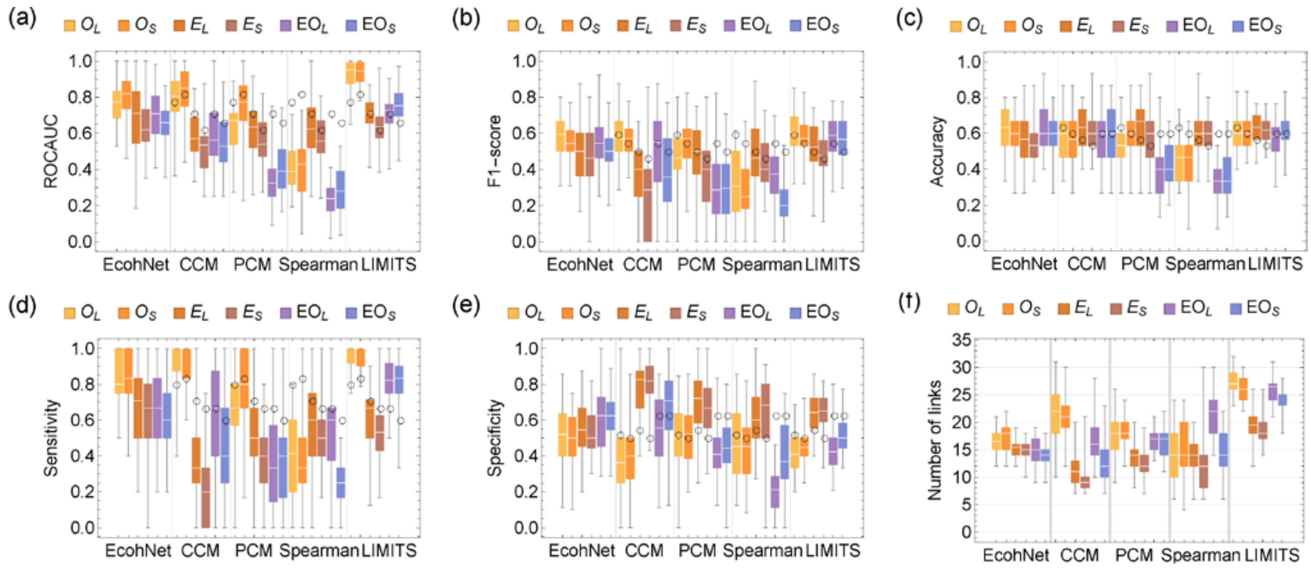

Figure S11. The value of ROC-AUC (a), F1-score (b), accuracy (c), sensitivity (d), specificity (e), and the number of inferred links (f) for the food web models with two unobserved species (8,000 steps sampled with an interval of 40 steps). In the box plot, white lines indicate the median, box edges indicate the first and third quartile value, and whiskers indicate maximum and minimum values. Black circles indicate the median value of EcohNet.

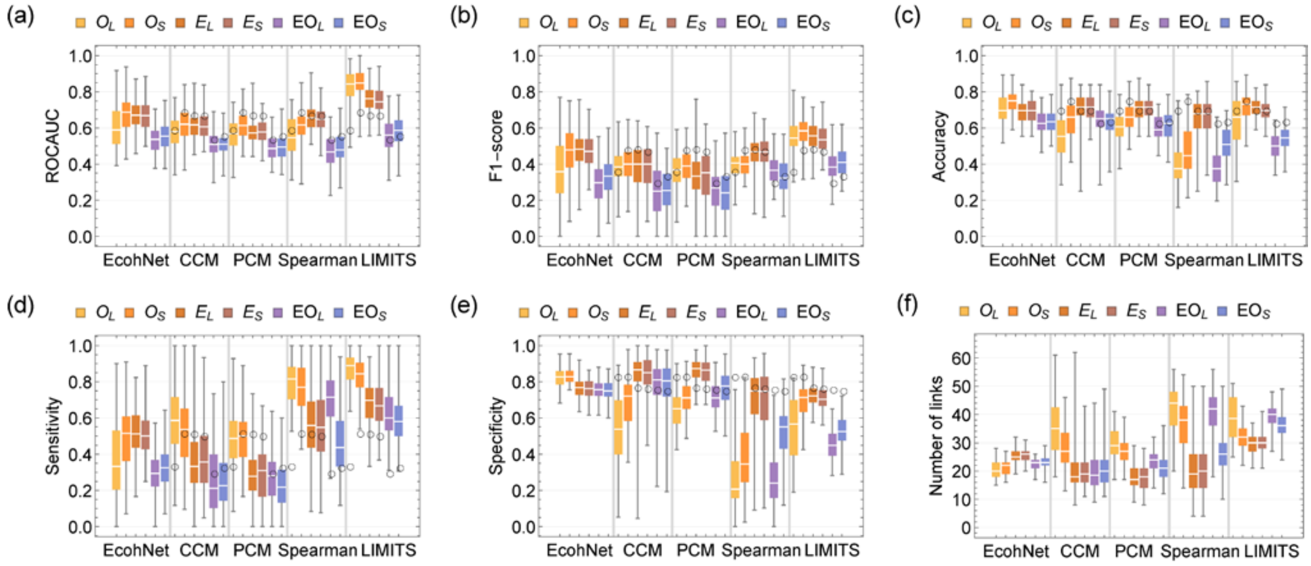

Figure S12. The value of ROC-AUC (a), F1-score (b), accuracy (c), sensitivity (d), specificity (e), and the number of inferred links (f) for the random interaction models. In the box plot, white lines indicate the median, box edges indicate the first and third quartile value, and whiskers indicate maximum and minimum values. Black circles indicate the median value of EcohNet. Relative to the food web models, in the random interaction model we considered only the ability of EcohNet, CCM and PCM to detect direct influences; we tested the presence of influence from one species to another (corresponding to a non-zero $a_{ij}$ in the interaction matrix) when there are causal links from the former to the later.

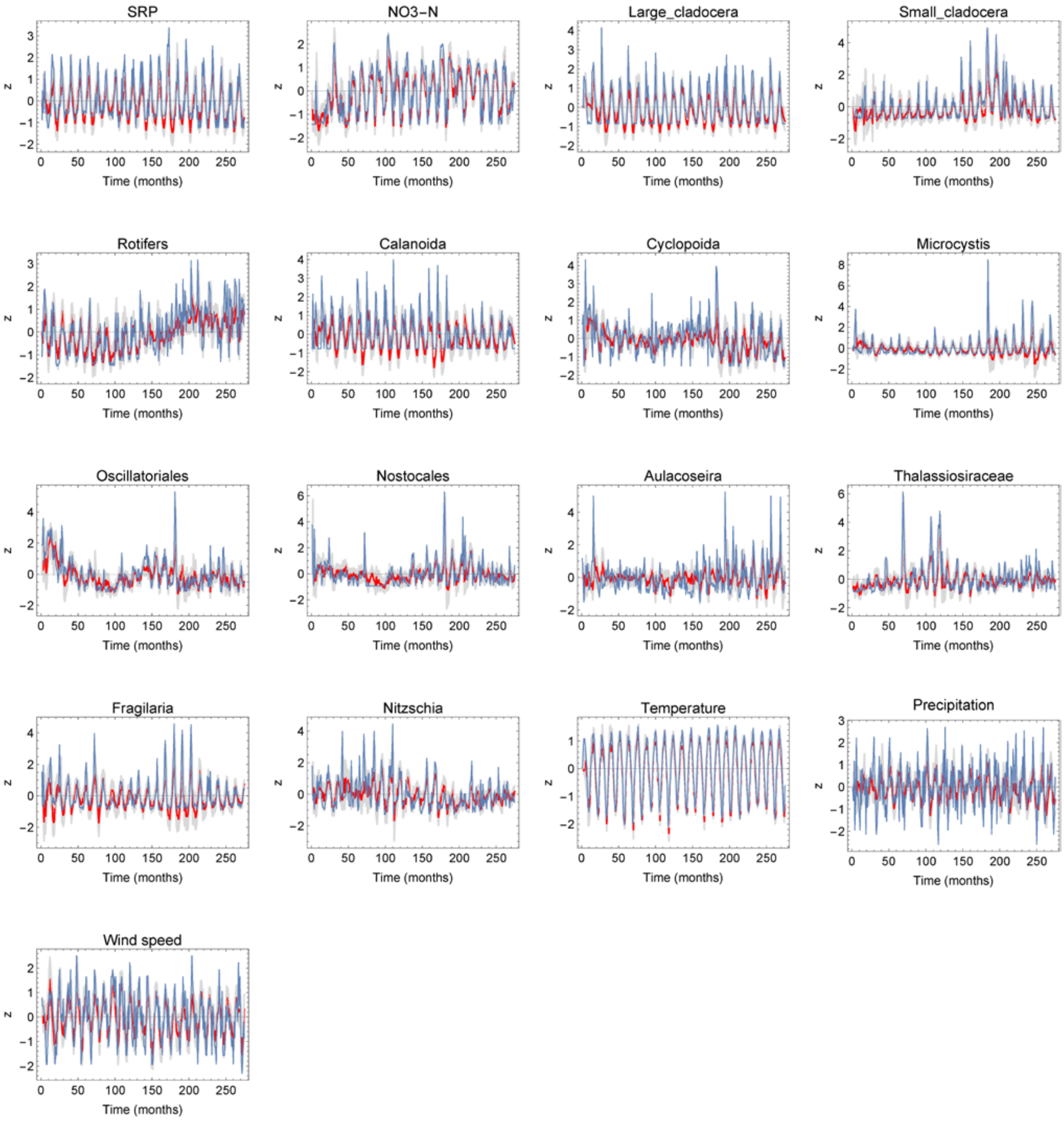

Figure S13. Results of prediction of each variables. Blue lines represent the actual time series and red lines represent the median of predicted time series with the 95% CI shown by grey areas (drawn by 10,000 time series predicted by EcohNet with optimal variables).

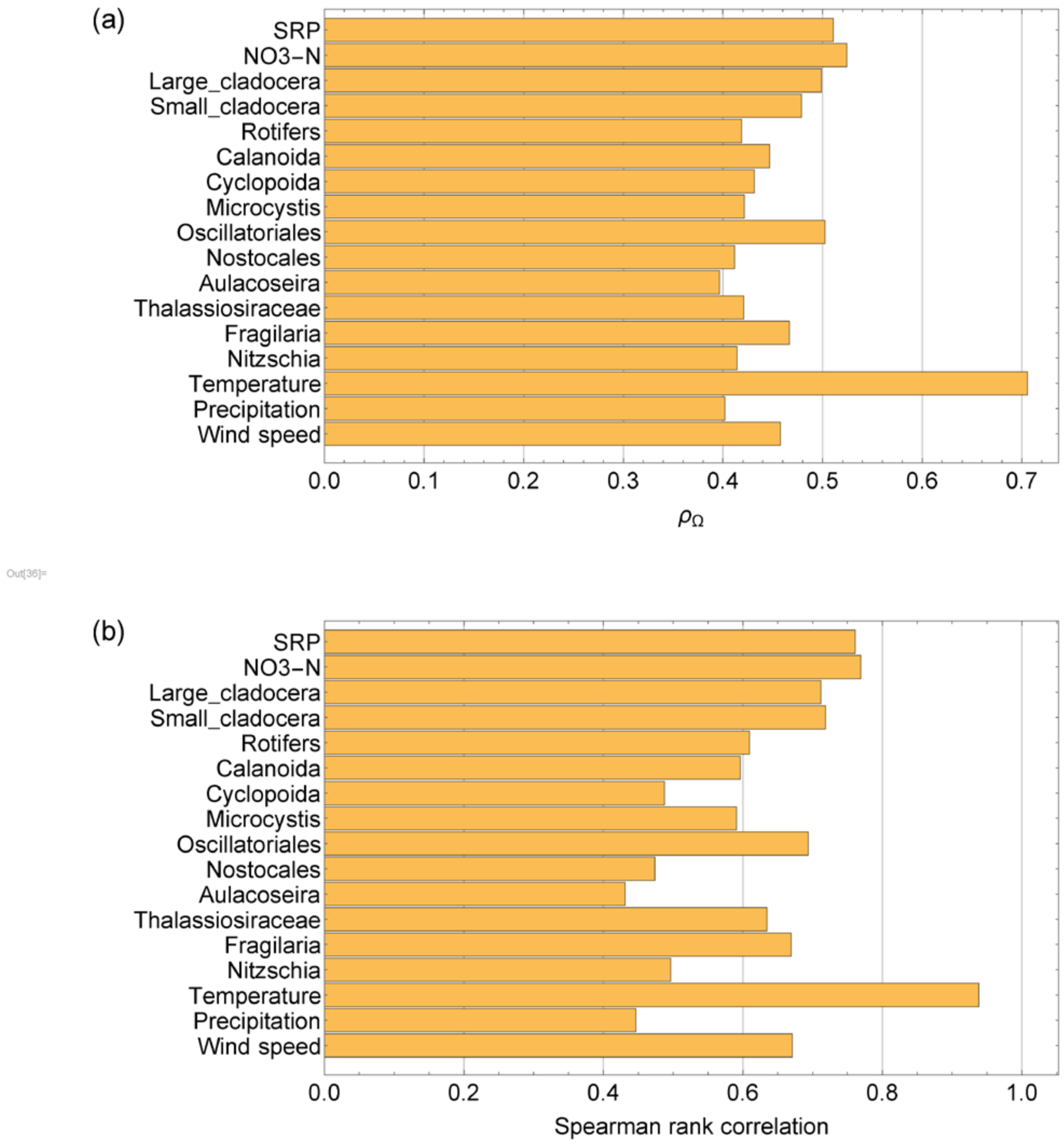

Figure S14. Performance of forecasting by EcohNet with the optimized variables. (a) Prediction skill of the median of forecasting by EcohNet. (b) Spearman rank correlation between the actual time series and the median of forecasting.

### SI Appendix Tables

Table S1. Summary of simulation conditions. For all conditions, 100 time series were generated for each of E, O, and EO, with two noise magnitudes S (0.05) and L (0.15), i.e., a total of 600 data sets were generated. Numerical simulations were performed with 16 species. Except for condition H, time series of 8 major species excluding 8 minor species were accepted. Condition H was generated by resampling 100 data sets from condition A with replacement and randomly excluding two species at that time. Numerical simulation results with lengths ranging from 2,000 to 16,000 (after discarding the first 2,000 steps) were sampled at intervals of 20 to 80 steps to generate data containing 50-200 data points; A-C have different numbers of data points; A, D-G have different sampling intervals. The model other than I is a food web model in which species are bi-directionally coupled by prey-predator links. Condition I is based on a random interaction model where species are coupled by unidirectional links. We used condition A as the representative condition.

| | A | B | C | D | E | F | G | H | I |
|---|---|---|---|---|---|---|---|---|---|
| Data size | 100×6 | 100×6 | 100×6 | 100×6 | 100×6 | 100×6 | 100×6 | 100×6 | 100×6 |
| Interaction type | food web | food web | food web | food web | food web | food web | food web | food web | random |
| Number of species | 8 (16) | 8 (16) | 8 (16) | 8 (16) | 8 (16) | 8 (16) | 8 (16) | 6 (16) | 8 (16) |
| Time series length | 8000 | 2000 | 4000 | 2000 | 4000 | 12000 | 16000 | 8000 | 8000 |
| Sampling interval | 40 | 40 | 40 | 10 | 20 | 60 | 80 | 40 | 40 |
| Number of data points | 200 | 50 | 100 | 200 | 200 | 200 | 200 | 200 | 200 |
| Unobserved species | minor species | minor species | minor species | minor species | minor species | minor species | minor species | minor species, two major species | minor species |
| Figure | Fig.2(g-j), 3, S16-S18 | Fig.S5 | Fig.S6 | Fig.S7 | Fig.S8 | Fig.S9 | Fig.S10 | Fig.S11 | Fig.S12 |

### A. Conventional approaches used for comparison

#### 1. *Spearman Correlation Coefficient*

Spearman rank correlation coefficient [1] is a nonparametric measure of rank correlation. It is relevant even if both or one of the variables is non-Gaussian and thus is more broadly applicable than Pearson correlation. The correlation coefficients are frequently used for network-based analysis of biological systems [2]. This method provides correlation coefficients between −1 and 1 for each pair of variables, accompanied by the p-values. The correlation matrix and p-values are symmetric.

#### 2. *Convergent Cross Mapping (CCM)*

CCM [3] is a statistical test for a causal relationship between two time series variables and attempts to address the problem that correlation is often not an indicator of the presence or absence of actual causal relationships. This method is based on Takens' embedding theorem [4], which states that the essential information of a multi-dimensional dynamical system is retained in the time series of any single variable of that system. Based on this theory, in a pair of time series $(X, Y)$, if $Y$ has a high forecasting skill in predicting $X$, causality will be detected in the direction of $X \rightarrow Y$. The predictive skill of CCM, denoted as $\rho^{ccm}_{X\rightarrow Y}$, is quantified by the Pearson correlation coefficient between actual $X$ and $X$ predicted by $Y$. Different from the correlation-based methods, $\rho^{ccm}_{X\rightarrow Y}$ is usually unequal to $\rho^{ccm}_{Y\rightarrow X}$.

To perform CCM, it is necessary to determine the appropriate embedding dimension for the state space reconstruction. We used a one-step-ahead self-prediction [3] to determine the embedding dimension $E$; we selected $E$ that maximizes prediction skill in the range $E = 1$ to $12$. Then, the strength of the causal relationship between X and Y was obtained as follows. First, we prepared the state space $M_Y$ reconstructed by the Y's embedding dimension $E_Y$. Then, in a leave-one-out manner, we predict X from $M_Y$ using the simplex projection method [3], which is labeled as $X^Y$. The cross-map prediction skill $\rho^{ccm}_{X\rightarrow Y}$ is the Pearson's correlation coefficient between $X$ and $X^Y$. To evaluate the significance of CCM we must test if cross-mapping skill improves with increasing library size [5]. There are several different approaches for the convergence test. We tested three representative approaches, namely, 1) with a surrogate time series with random sampling of reconstructed state spaces, 2) with a surrogate time series with random sampling of fixed intervals in the time series, 3) without a surrogate time series with random sampling of points in the reconstructed state, and used 1) throughout this paper as it performed best in the benchmark using a food-web simulation (Fig. S15).

1) As a first approach, we followed the implementation by Clark et al. [6]; we calculated $\rho_{max} = \rho^{ccm}_{X\rightarrow Y}$ with the maximum library length (i.e., full-length of available time series) as the prediction skill representing causality from X to Y. We then sampled $E + 1$ of points in $M_Y$ constructed with the full-length of Y without replacement and repeated the same procedure as above to obtain the prediction skill. We repeated this procedure $N = 500$ times to obtain a distribution of prediction skills $F_{min}$. We also obtained another set of prediction skills, $F_{surrogate}$ without sampling, but with a surrogate time series. To

obtain a surrogate time series, the full-length of Y was shuffled by the Ebisuzaki method [3, 7] when the target system has no explicit periodicity (aquatic mesocosm data, $O$ and $E$). The Ebisuzaki method randomizes a time series while retaining its power spectrum. When the target system is forced by a periodic oscillation ($EO$, Kasumigaura monitoring data) the seasonal surrogate method [8] is used. In the seasonal surrogate method, the phase average $\tilde{X}(t) = \tilde{X}(t+T)$ (e.g., for a seasonal cycle with cycle $T$) and the anomaly as the difference between the original time series and the phase average $X^*(t) = X(t) - \tilde{X}(t)$ was calculated. Then $X^*(t)$ was randomly shuffled, and added back to $\tilde{X}(t)$ to obtain a surrogate time series. P-value was calculated as $max(p_1, p_2)$ (this is not a p-value in the strict sense, but it aims to prevent the matrix from becoming extremely dense) where $p_1$ and $p_2$ is the ratio of elements in $F_{min}$ and $F_{surrogate}$, respectively, that are larger than $\rho_{max}$.

2) The second approach is the same as the first approach except that when obtaining $F_{min}$, the length of the $2E$ interval (corresponding to $E+1$ points in the reconstructed phase space) was randomly sampled from the full-length of Y. This sampling scheme is based on the original implementation of Sugihara et al. [3] and adopted in many other studies.

3) In the third approach, instead of calculating the surrogate time series, the distribution of prediction skills was obtained with four different samplings, which was 2E+1, 25%, 50% and 75% of points in $M_Y$ constructed with the full-length of Y. These points were then used to calculate Kendall $\tau$ to test convergence. Instead of the p-value calculated by $F_{surrogate}$, p-value of Kendall $\tau$ was used as $p_2$. This approach was used in several recent studies [5, 9, 10].

3. *Partial Cross Mapping (PCM)*

PCM [11] is an extension of CCM that incorporates partial correlation. The key idea is to examine the consensus between one time series and its cross-map prediction from the other with conditioning on the part that is transferred from the third variable. For the chain relationship X←Z←Y, the direct causality between X and Y is calculated as $\rho^{pcm}_{X\to Y|Z} = |PCC(X, X^Y|X^{Z^Y})|$. Here, $X^{Z^Y}$ is obtained by a successive simplex projection ($X^{Z^Y}$ is X predicted by $M_{Z^Y}$ where $Z^Y$ is Z predicted by $M_Y$) and characterizes the indirect information flow between X and Y through Z. $PCC(x,y|z) = \frac{Corr(x,y) - Corr(x,z)Corr(y,z)}{\sqrt{(1-Corr(x,z)^2)(1-Corr(y,z)^2)}}$ is the partial correlation coefficient describing the association degree between $x$ and $y$ with information about the $z$ removed. Thus, $\rho^{pcm}_{X\to Y|Z}$ is the direct causation from X to Y conditioned on the indirect causation through Z. The above definition can be extended to a multivariate relationship as, $\rho^{pcm}_{X\to Y|\{Z_i\}} = |PCC(X, X^Y|\{X^{Z_i^Y}|i = 1,2,\dots\})|$ by recursively applying the relationship, $PCC(X,Y|Z_1,Z_2) = \frac{PCC(X,Y|Z_1) - PCC(X,Z_2|Z_1)PCC(Y,Z_2|Z_1)}{\sqrt{(1-PCC(X,Z_2|Z_1)^2)(1-PCC(Y,Z_2|Z_1)^2)}}$. Following the original implementation of the authors, we used the false neighborhood method to calculate the embedding dimension for reconstructed state space $M$; we selected $E$ in the range $E = 1$ to $12$.

4. *LIMITS*

LIMITS [12] is based on the generalized Lotka–Volterra difference equation. The algorithm combines forward stepwise regression and bootstrap aggregation (bagging) to determine, in a majority voting fashion, the pairs of interacting variables and the value of their interaction coefficient. For each species, the logarithm of the change in abundance per time step is used as the response variable. Then, this method selects variables to be explanatory variables of linear regression step by step, as long as the predictive performance is improved (more precisely, a threshold level of the improvement of prediction error is need to be set to stop adding a new variable when it exceeded). The species selected as the explanatory variables are candidates for interaction partners. This process is repeated (here, 256 times) as bootstrap replicates, and the species selected in more than half of all iterations are finally determined as the interaction partners, i.e., variables that can affect the response variable. The interaction is estimated asymmetrically and in a way that includes positive and negative values. This method returns a sparse matrix for the interaction matrix.

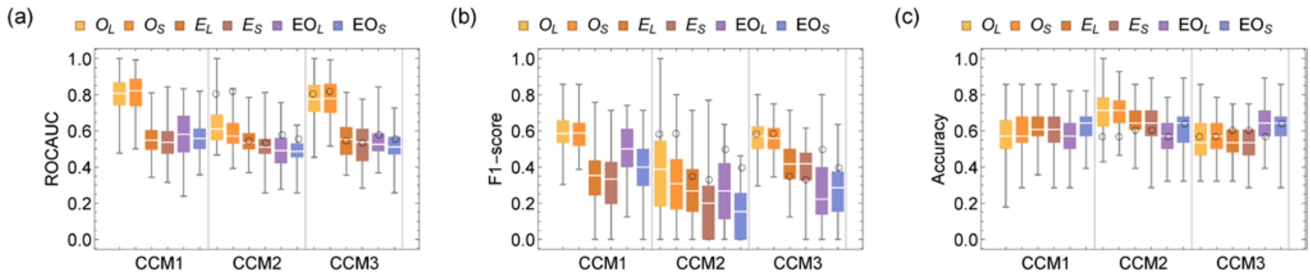

Figure S15. The value of ROC-AUC (a), F1-score (b) and, accuracy (c) for food-web simulation (8,000 steps sampled with an interval of 40 steps). Three different implementations were tested: 1) with a surrogate time series with random sampling of reconstructed state spaces, 2) with a surrogate time series with random sampling of fixed intervals in a time series, 3) without a surrogate time series with random sampling of points in a reconstructed state. In the box plot, white lines indicate the median, box edges indicate the first and third quartile value, and whiskers indicate maximum and minimum values. Black circles indicate the median value of CCM1.

### B. Parameters and performance of ESN-RLS

The relationship of the predictive performance of ESN-RLS and the four representative parameters, spectral radius ($\rho_0$), forgetting factor ($\lambda$), regularization factor ($\delta$), and the number of ESN updates ($\tau$) is as follows. As $\rho_0$ becomes smaller than 1, the ability of ESN to track a time series becomes lower and it cannot adequately deal with complex time series. On the other hand, when $\rho_0$ becomes larger than 1, the behavior of ESN becomes unstable since $\rho_0 < 1$ ensures echo state property in most situations [16]. Thus, it is recommended that $\rho_0$ be set to a value close to 1, but smaller than 1. $\lambda$ determines how fast the effects of past errors decay. The closer the value is to 1, the slower the decay, and at 1, all past information is treated with the same weight. It is also recommended that $\lambda$ be set to a value smaller than 1 and close to 1 [13]. $\delta$ controls the strength of the normalization in the regression and it is recommended to set $\delta \ll 1$ [15]. Since $\tau$ affects how well an ESN can respond to changes in the short time series, its effect is complementary to $\rho_0$ and $\lambda$. To keep the basic flexibility high enough, $\tau$ would need to be greater than 1.

The performance of EcohNet is essentially related to the performance of ESN-RLSs to track time series. Our validation (Fig. 16S) showed that the performance of EcohNet is best in the range where ESN-RLSs are generally balanced in tracking time series and stabilizing dynamical behavior, and it is consistent with the general recommendations above. We actually found that $\tau$ need to be sufficiently greater than 1, and that $\rho_0$ and $\lambda$ must be close to 1. The evaluation criteria did not vary significantly for small differences in parameters except when $\lambda$ approached 1; in this case, since the decay time is scaled by $(1-\lambda)^{-1}$, the closer $\lambda$ is to 1, smaller differences in $\lambda$ result in larger differences in the decay time.

There is little concern about parameter tuning for ESN-RLS. In fact, even without changing the parameters chosen here ($(\rho_0, \lambda, \delta, \tau) = (0.95, 0.95,\ 0.001, 8)$), EcohNet outperformed the other methods in the evaluation on simulated data.

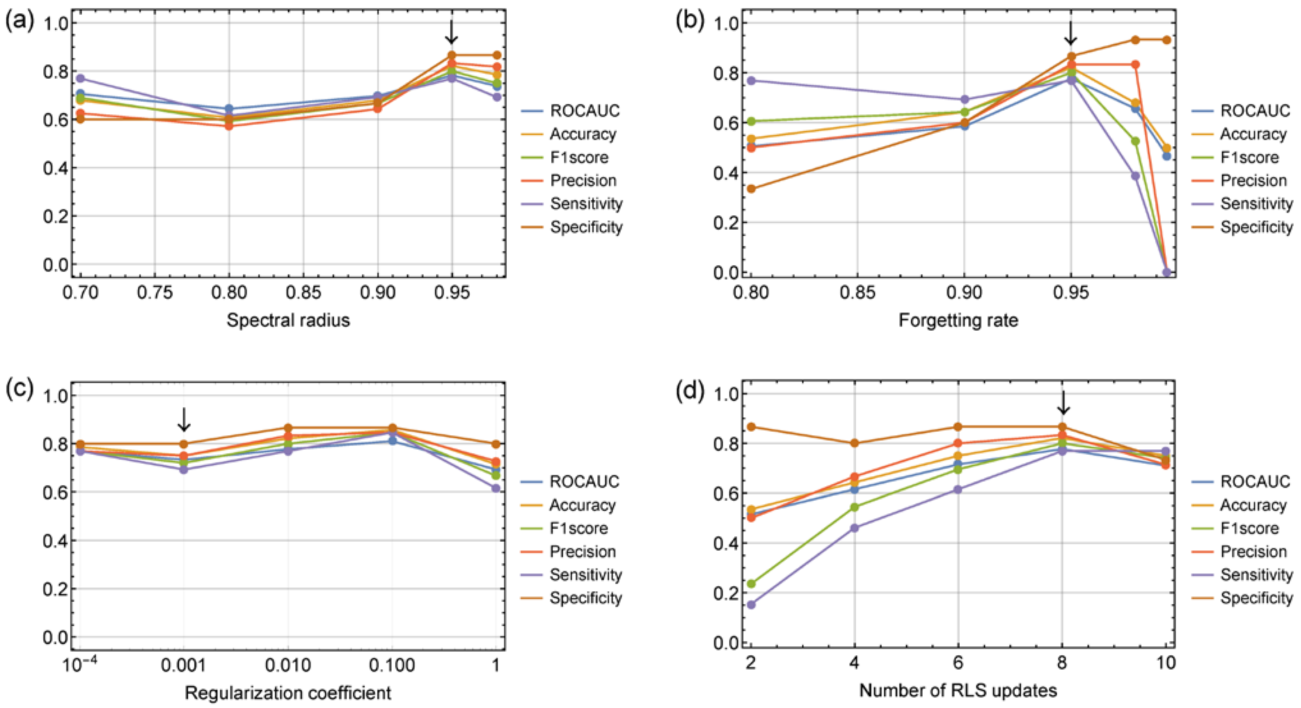

Figure S16. Sensitivity of evaluation criteria for the ESN-RLS parameters. (a) spectral radius ($\rho_0$), (b) Forgetting factor ($\lambda$), (c) Regularization coefficient ($\delta$), and (d) Number of RLS updates ($\tau$). The parameter values used in this study are marked with black arrows.

### C. Evaluating the ability to detect direct influences in food web models

The benchmarks using the mesocosm experiment and food web simulation data in this paper considered the ability of EcohNet, CCM and PCM to detect interacting pairs. When evaluating the ability to detect direct influences from one species to another (as in the random interaction models), the advantage of EcohNet over CCM appeared to be reduced in the food web models (Fig. S17). Specifically, CCM outperformed EcohNet in ROC-AUC for $O_{S,L}$ and $O_{S,L}$ and $EO_L$ for F1-score. However, this superiority of CCM was probably due to two factors: 1) the interaction matrix of a food web model is symmetric, and 2) CCM tends to detect causality in a symmetric manner. In fact, the ratio of symmetric causal links (causal links detected both $X \rightarrow Y$ and $Y \rightarrow X$) to the total number of causal links remained the same in CCM both in the food web models and the random interaction models, and the value was larger than EcohNet (Fig. S18). This estimation bias could result in the superiority of CCM when correspondence of the directionality of causal links was considered in the food web models. In fact, EcohNet outperformed CCM in ROC-AUC and F1-score in the random interaction models in which interaction matrices were asymmetric and ability to detect the direct influence through the causal links was considered.

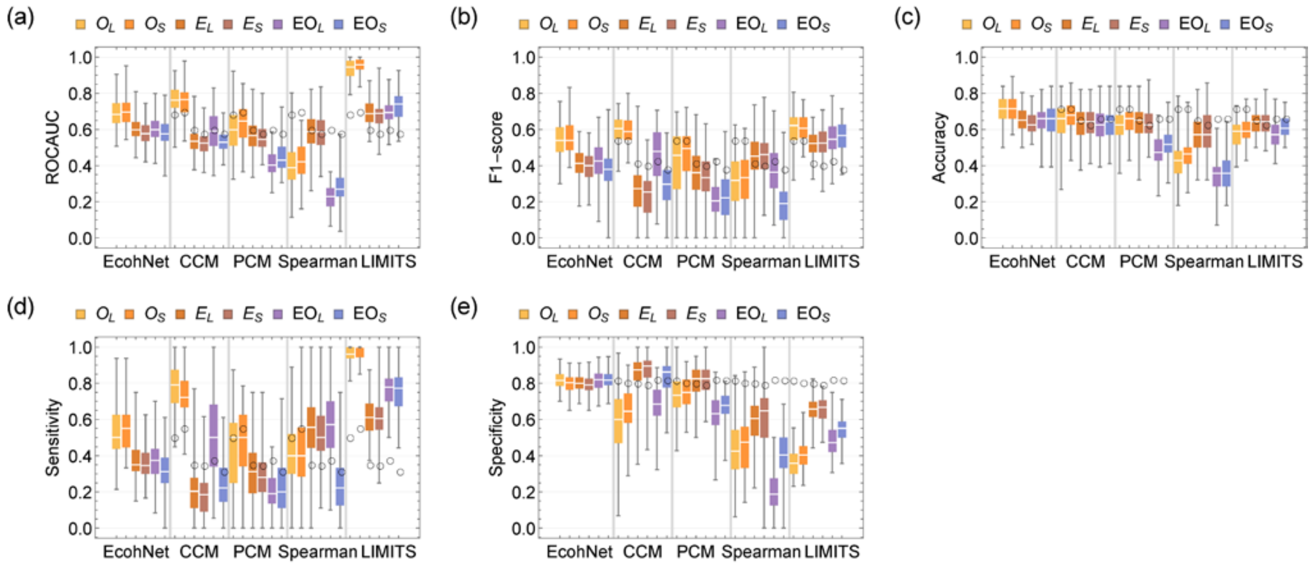

Figure S17. The value of ROC-AUC (a), F1-score (b), accuracy (c), sensitivity (d), and specificity (e) for the food web models (8,000 steps sampled with an interval of 40 steps) when the directionality of causal links was considered in EcohNet, CCM and PCM. We tested the presence of influence from one species to another (corresponding to a non-zero $a_{ij}$ in the interaction matrix) when there were causal links from the former to the later. In the box plot, white lines indicate the median, box edges indicate the first and third quartile value, and whiskers indicate maximum and minimum values. Black circles indicate the median value of EcohNet.

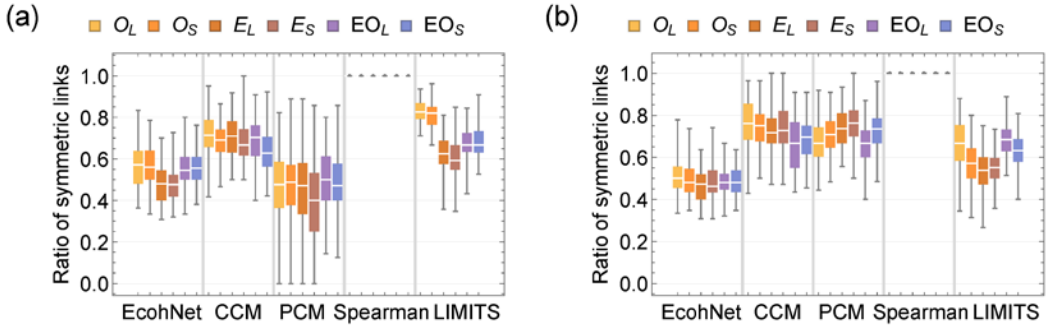

Figure S18. Ratio of symmetric links to the total number of links in the food web models (8,000 steps sampled with an interval of 40 steps) (a) and the random interaction models (b). In EcohNet, CCM and PCM, the ratio of symmetric causal links to the total number of causal links was considered, and in LIMITS, the ratio of symmetric interactions to the total number of interactions was considered.

### D. Description of datasets

#### 1. *Long-term mesocosm experiment*

We first benchmarked our method using the dataset (transformed_data_Nature2008) of a long-term mesocosm experiment by Beninca et al. [17], which is available as a supplement of Beninca et al. [18] (https://onlinelibrary.wiley.com/action/downloadSupplement?doi=10.1111%2Fj.1461-0248.2009.01391.x&file=ELE_1391_sm_appendix+S1.xls). The mesocosm consisted of a cylindrical plastic container (74cm high, 45cm diameter), which was filled with a 10cm sediment layer and 90 $\ell$ of water from the Baltic Sea. The mesocosm was maintained in the laboratory at a temperature of about 20℃, a salinity of about 9‰, incident irradiation of 50 $\mu$ mol photons $m^{-1}s^{-1}$ (16:8 hour light: dark cycle) and constant aeration.

The authors identified three different strengths of interactions between species in this mesocosm. We included all of them to evaluate the performance of our selected methods. However, we only considered 9 components (Calanoids, Rotifers, Ostracods, Nanophytoplankton, Picophytoplankton, Bacteria, Total dissolved inorganic nitrogen, Solvable reactive phosphorus (SRP)) to use a time series where as many species as possible coexisted. As a result, we used data from 700 days (210 points) out of the 2319 days of the observation period. We set the parameters of the ESN-RLS according to the results of the sensitivity analysis for this data (Fig. S16) and kept it fixed throughout this paper.

#### 2. *Food web simulation*

We used a food web model to generate a data set for benchmarking. The model is based on a generalized Lotka-Volterra equation (GLVE),

$$\frac{dx_i}{dt} = F_i(x) = x_i\left\{r_i\left(1 - \frac{\Sigma_{j=1}^{M} x_j}{K}\right) + e^{-1}\Sigma_{j=1}^{M}\frac{a_{ji}x_i x_j}{\beta_i + \Sigma_{k=1}^{M} a_{ki}x_k} - \Sigma_{j=1}^{M}\frac{a_{ij}x_i x_j}{\beta_j + \Sigma_{k=1}^{M} a_{kj}x_k} - mx_i\right\}, \quad (1)$$

which has been frequently used to benchmark methods for inferring interaction networks [12, 19, 20]. Here, $x_i$ is the abundance of species $i$, $r_i$ is the intrinsic growth rate, $K$ is the carrying capacity, $e$ is the conversion efficiency, $a_{ij} \in A$ is the per capita grazing effect of species $j$ on $i$ (matrix $A$ is an interaction matrix that represents the strength of interaction between species and is symmetric), $\beta_i \in \beta$ is the half-saturation constant of the interspecific interaction, and $m$ is the mortality rate. $M$ is the number of species and we set $M = 16$ throughout this paper.

Based on eq. (1), we generated time series by numerically solving,

$$x_i(t + \Delta t) = x_i(t) + F(x)\Delta t + \eta_i(t) + \epsilon. \quad (2)$$

Here, $\eta_i(t)$ is a random vector whose elements are drawn from a normal distribution with mean 0 and standard deviation $\sigma\sqrt{\Delta t}$, $\Delta t$ is the time step for numerical simulation, and $\epsilon$ is the small size of influx introduced to avoid underflow. We solved eq.(2) for 4,000 to 18,000 steps (SI Appendix Table S1) with $\Delta t = 0.05$ and dropped the first 2,000 steps to remove the transient. Then, we resampled the time series with an interval of 10 to 80 and obtained a time series with 50, 100, and 200 points (SI Appendix Table S1). We used a case with 8,000 steps sampled with an interval of 40 steps (200 points) as the representative result. Each data set for different conditions consists of 100 time series simulated with randomly generated $A$ and $\beta$. For a data set, a time series is accepted if the number of major species (here, it is defined as a species whose mean abundance is larger than 0.01) is equal to 8, and the abundance of species other than the major species is removed. Thus, one time point of a time series in a data set is a vector of eight elements. This corresponds to the fact that some of the minor species may not be included in the observations. Moreover, to evaluate the effect of missing major species, we considered a condition where two major species were removed from the original time series (Table S1). Before the analysis, the time series were normalized to have a mean 0 variance of 1.

To generate the time series of different dynamical complexity, we used three different parameter settings and two different noise magnitudes. The parameter settings are represented as $E$, $EO$ and $O$. $E$ is the equilibrium in which $e = 0.5$ and $m = 2$, $EO$ is the equilibrium forced by an external oscillator in which $e = 0.5$ and $m = 2$ but K is replaced by a function $K(t) = 1 + 0.8 \sin t/20\pi$, $O$ exhibits intrinsic oscillation with $e = 0.33$ and $m = 1$. We used two different values $\sigma = 0.05$ and $0.15$ for small and large noise magnitude, and indicated them by suffix S and L, e.g., $E_S$ and $E_L$. Other parameter values were as follows. The interaction matrix specifies the food web by a cascade model [21]. Here, if the species' index $j$ is greater than $i$, $j$ preys on $i$ with probability 0.33 ($a_{ij} = 0$ if $i \geq j$ and otherwise randomly drawn from a uniform distribution between $0.05$ and $2$ with probability 0.33 and otherwise 0). We set $r = 1$ and $K = 1$ for the species having no prey and $r = -1$ and $K = 10^8$ otherwise. The value of $\beta$ was randomly assigned between $0.01$ and $0.1$.

3. *Random interaction model*

The random interaction model is given as the following equation:

$$\frac{dx_i}{dt} = x_i \left\{ r_i \left(1 - \frac{b \Sigma_{j=1}^{M} a_{ji} x_j}{K}\right) - m x_i \right\}$$

In this model, species interactions are defined only by negative influences (i.e., $a_{ij} < 0$). As in the case of the food web model, the connectance of interaction matrix $A$ was set to 0.33 but the non-zero elements were selected randomly from the off-diagonal part. Thus, $A$ is an asymmetric matrix. Other parameters were set according to the food web model except that the parameter settings corresponding to $E$, $EO$ and $O$ were $b = 2$ and $m = 2$, $b = 2$ and $m = 2$ and $b = 4$ and $m = 1$, respectively.
